# Loss of *Bicra/Gltscr1* leads to a defect in fetal liver macrophages responsible for erythrocyte maturation in mice

**DOI:** 10.1101/2024.10.17.618940

**Authors:** Surbhi Sood, Aktan Alpsoy, Guanming Jiao, Alisha Dhiman, Charles Samuel King, Gabriella Grace Conjelko, Judy E. Hallett, Sagar M Utturkar, Jill E Hutchcroft, Emily C Dykhuizen

**Affiliations:** Borch Department of Medicinal Chemistry and Molecular Pharmacology, Purdue University, West Lafayette, Indiana, USA; Purdue Institute for Cancer Research, Purdue University, West Lafayette, Indiana, USA; Purdue University Interdisciplinary Life Science Graduate Program (PULSe); Bindley Bioscience Center, Purdue University, West Lafayette, Indiana, USA

## Abstract

**Key points:** *Bicra/Gltscr1* homozygous knockout mice are perinatal lethal with aberrant liver resident macrophage gene expression and function.

Dysfunctional macrophages result in accumulation of immature nucleated red blood cells in peripheral blood and liver of the knockout mice.

GLTSCR1, a protein encoded by the *Bicra* gene, is a defining subunit of the SWI/SNF (also called mammalian BAF) chromatin remodeling subcomplex called GBAF/ncBAF. To determine the role of GLTSCR1 during mouse development, we generated a *Bicra* germline knockout mouse using CRISPR/Cas9. Mice with homozygous loss of *Bicra* were born at Mendelian ratios but were small, pale and died within 24 hours after birth. Histology indicated blood-related defects including defective erythroblastic islands and irregularly sized red blood cells. Gene expression profiling of fetal livers pinpointed a defect in liver resident macrophages involved in the last stage of erythrocyte maturation, resulting in accumulation of nucleated erythrocytes in *Bicra^-/-^* pups. Together, these results demonstrate that *Bicra* is critical for fetal liver macrophage function during development.

Visual Abstract

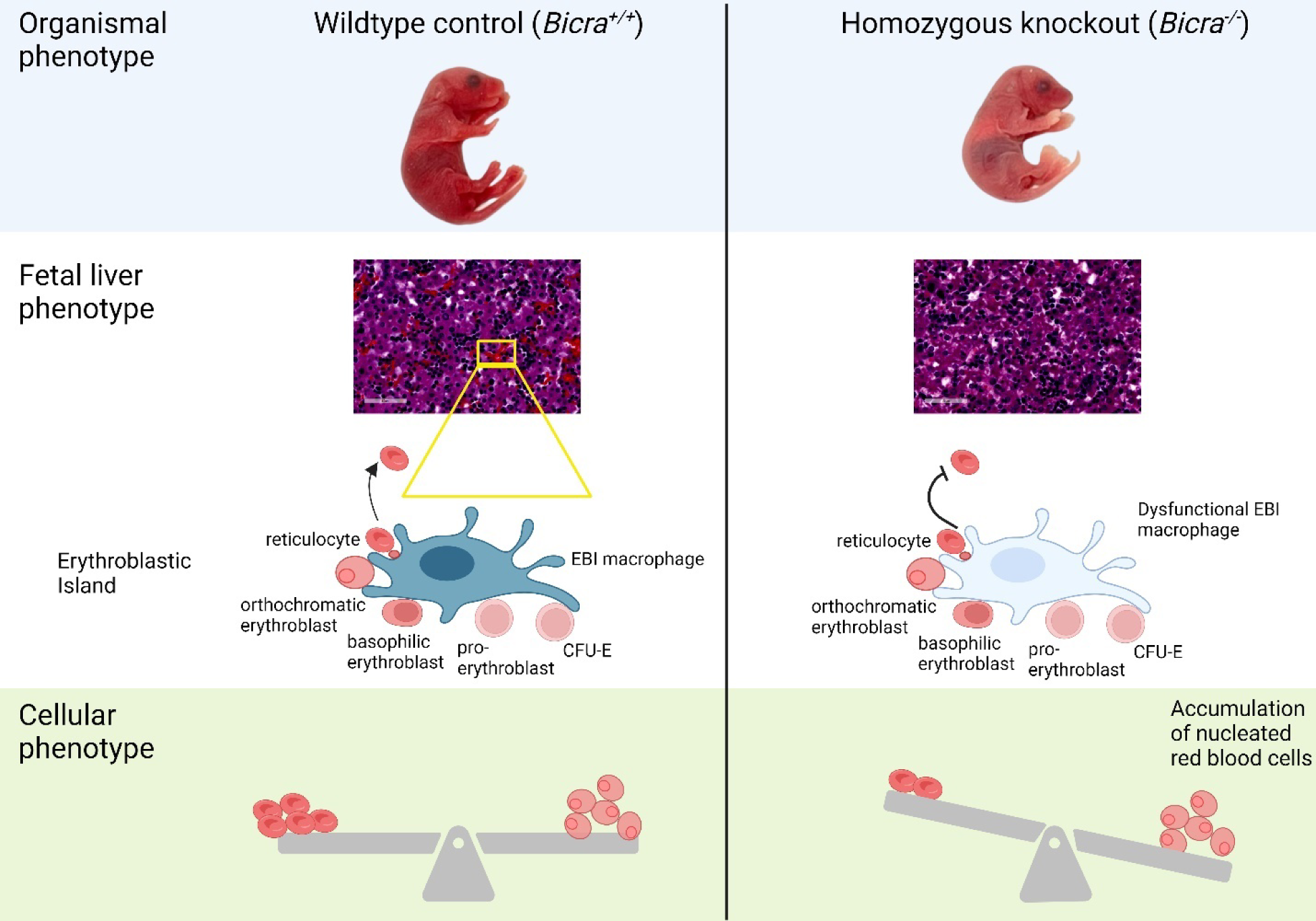

## Introduction

In eukaryotic cells, DNA is packaged into chromatin (Luger et al. 1997). The fundamental repeating unit of chromatin is the nucleosome which is composed of a histone protein octamer wound by DNA (Kornberg 1974). This chromatin structure is highly dynamic and is altered during embryonic development to facilitate tissue and cell type specification (Woodcock and Ghosh 2010). Misregulation of chromatin results in aberrant downstream gene expression (Berger 2007) and developmental defects (Bickmore and van Steensel 2013).

The mammalian <u>Swi</u>tch/<u>s</u>ucrose <u>n</u>on-fermenting (mSWI/SNF) chromatin remodeling complex, also known as the <u>B</u>RG1<u>-a</u>ssociated factors (BAF) complex (W. Wang et al. 1996), utilizes ATP to reposition nucleosomes to modulate DNA accessibility (Imbalzano et al. 1994; Gutiérrez et al. 2007; Cairns 2007) and gene expression during development (Ho and Crabtree 2010). mSWI/SNF can exist in three biochemically distinct subcomplexes: canonical BAF (cBAF), Polybromo1 containing BAF (PBAF) and GLTSCR1/1L containing BAF (GBAF, also referred to as non-canonical/ncBAF) (Alpsoy and Dykhuizen 2018; Gatchalian et al. 2018; Mashtalir et al. 2018). GBAF uniquely contains BRD9 and one of the two mutually exclusive paralogous subunits GLTSCR1/*BICRA* or GLTSCR1L/*BICRAL*. Specific inhibitors and degraders of BRD9 are commonly used for assessing GBAF function in dependent cell lines (Michel et al. 2018; Brien et al. 2018; Weisberg et al. 2022; Kurata et al. 2023; X. Wang et al. 2019). Even in BRD9-dependent cells, GLTSCR1 and GLTSCR1L are often co-expressed and functionally redundant (Alpsoy and Dykhuizen 2018; Gatchalian et al. 2018). Unique biological functions for these two paralogs remain undefined.

To identify non-redundant functions for *Bicra in vivo*, we used CRISPR-Cas9 gene editing to generate a germline knockout mouse. Mice with homozygous deletion of *Bicra* were born at Mendelian ratios but died shortly after birth with anemia and dyspnea. Through a combination of phenotypic and functional assays, we describe here an essential role for *Bicra* in the function of liver resident macrophages involved in terminal erythrocyte maturation.

## Results

### Generation of Bicra knockout mice and its phenotypic characterization

The pronuclei of fertilized mouse eggs from C57BL/6J (CD45.2^+^) mice were injected with mRNA encoding Cas9 and a single guide RNA (sgRNA) construct targeting exon 4 of *Bicra*, which was previously validated in mouse ES cells (Alpsoy and Dykhuizen 2018) (**Figure 1A**). Embryos were implanted in female C57BL/6J mice, which yielded 56 offspring. We genotyped the founder population (F_0_ offspring) by sequencing a window of 400 base pairs flanking the Cas9 ribonucleoprotein cut site on *Bicra* gene using Sanger sequencing (**Figure 1B**). Single allele editing was detected in 9 pups (8 male, 1 female). Using a male with 8bp deletion in one allele, we backcrossed the *Bicra^+/-^* males five times to wildtype C57BL/6J females.

**Figure 1:**
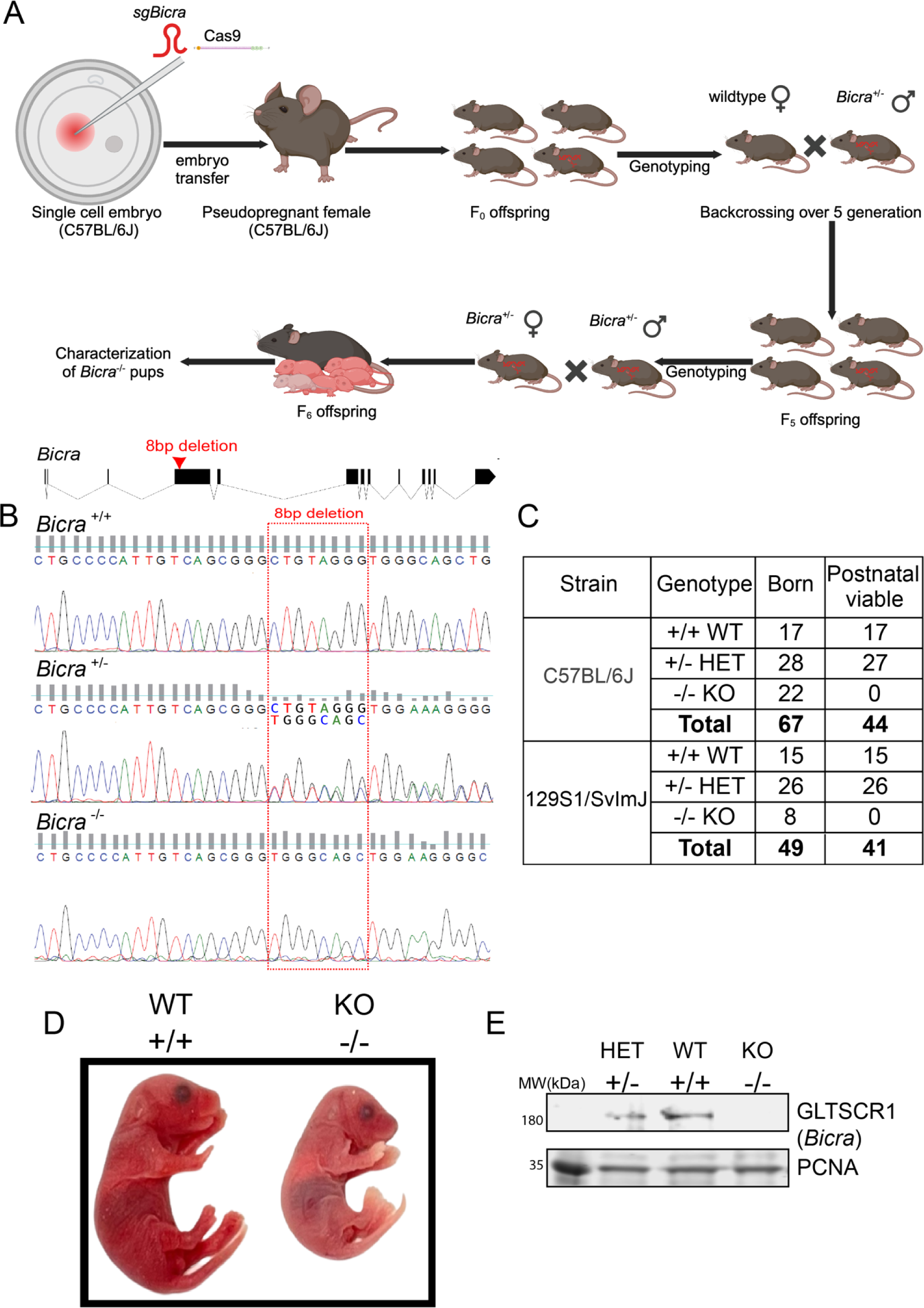
Generation of *Bicra* knockout mouse strain. (A) Schematic illustrating CRISPR/Cas9-mediated generation of *Bicra* knockout mice. (B) Exon diagram for *Bicra* gene and representative chromatogram from mouse of each genotype. (C) The genetic transmission Mendelian distribution table (Chi-square test; χ^2^ =2.552 (C57BL/6J), χ^2^ =1.137 (129S1/SvImJ) and df=2. Two-tailed P value is 0.2791) (D) Comparative morphology of *Bicra*^+/+^ (wildtype) and *Bicra*^-/-^ (homozygous knockout) C57BL/6J embryos at birth. (E) Protein lysates isolated from C57BL/6J livers at E14.5; PCNA is used as a loading control.

*Bicra*^+/-^ heterozygous mice were born at normal Mendelian ratios, were fertile, lived a normal lifespan, and were phenotypically indistinguishable from wild type littermates. Intercrossing of *Bicra^+/-^*mice yielded expected Mendelian ratios of *Bicra*^-/-^, *Bicra*^+/-^, and *Bicra*^+/+^ offspring at birth (**Table 1C**); however, *Bicra^-/-^*pups died shortly after birth. Most pups died within an hour of birth while some were able to survive several hours; none survived longer than 24 hours. We confirmed the loss of GLTSCR1 protein in E14.5 embryos (**Figure 1E**) and in mouse embryonic fibroblasts (MEFs) from E12.5 embryos (**Figure S1A**). To confirm that the phenotype is robust and strain-independent, we backcrossed male *Bicra*^+/-^ mice five times with wildtype 129S1/SvImJ females and observed a similar perinatal lethality for the *Bicra*^-/-^ pups (**Table 1C**). In both strains, the homozygous knockout pups were paler (**Figure 1D**) and had labored breathing compared to their littermates (**Supplementary video 1**), indicating a potential defect in heart, lung, or blood.

**Table 1:**

| Sr. No. | Antibody | Catalog | Vendor |
| --- | --- | --- | --- |
| 1. | Brilliant Violet 421™ anti-mouse CD71 | 113813 | BioLegend |
| 2. | PE/Cyanine5 anti-mouse CD45 | 103109 | BioLegend |
| 3. | PE anti-mouse TER-119 | 116208 | BioLegend |
| 4. | Alexa Fluor 647 anti-mouse F4/80 Antibody | 123121 | BioLegend |
| 5. | Pacific Blue anti-mouse/human CD11b | 101223 | BioLegend |

### Bicra^-/-^ knockout mice have morphological defects in peripheral erythrocytes

To investigate whether there are any obvious anatomical defects that might explain the anemia and perinatal lethality, we performed H&E staining on newborn *Bicra*^+/-^ and *Bicra*^-/-^ littermates (**Figure 2A)**. Gross histology of the *Bicra*^-/-^ pups pointed to a prominent defect in red blood cells in the liver, which is the main site of definitive erythropoiesis from E10.5 until birth when it switches to the bone marrow (Palis et al. 1999). The fetal liver is the major hematopoietic organ during development, supporting HSC (hematopoietic stem cell) expansion, active erythro-myeloid hematopoiesis, and erythrocyte maturation(Lewis, Yoshimoto, and Takebe 2021). The most prominent defect based on H&E staining was a significant reduction in mature erythroblastic islands (EBI) in the liver of *Bicra*^-/-^ pups at birth (**Figure 2B, 2D**) and at E14.5 (**Figure S2A, S2B and S2C**). EBI are specialized anatomic structures found in bone marrow, spleen and liver that facilitate proliferation and terminal maturation of red blood cells in association with a central macrophage (Manwani and Bieker 2008). In addition to the reduction of EBIs in the liver, the red blood cells in major vessels of the heart were rounder and paler in color compared to littermate controls (**Figure 2C, 2E**) In mammals, defects in primitive erythropoiesis, which occurs in the yolk sac, are embryonic lethal; however defects in definitive erythropoiesis, which occur in the fetal liver, are often associated with perinatal lethality and consistent with the phenotype of the *Bicra*^-/-^ pups (Baron, Isern, and Fraser 2012). Therefore, we next focused on defining the role of *Bicra* in definitive erythropoiesis in the fetal liver.

**Figure 2:**
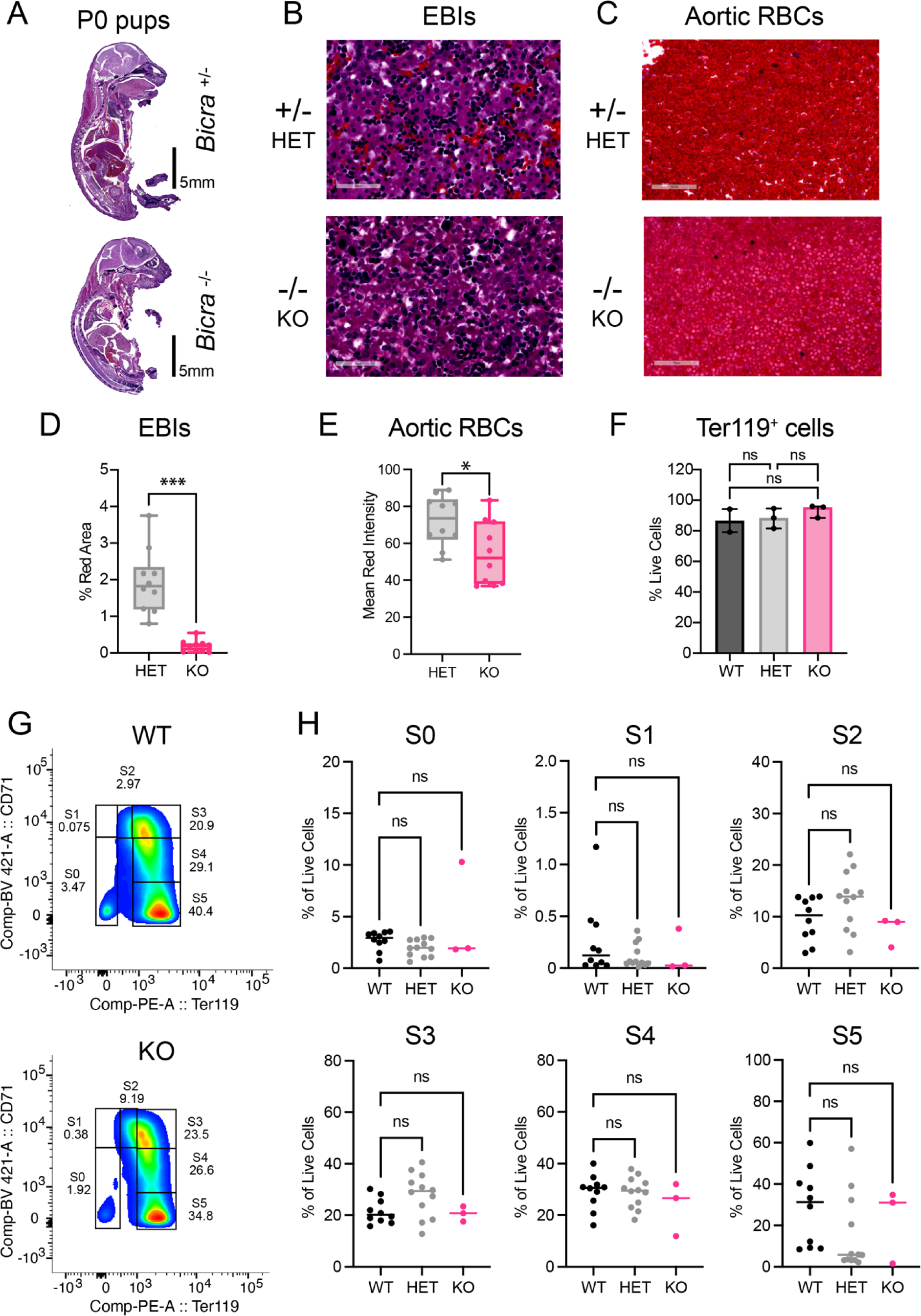
Histological characterization of *Bicra* knockout mouse strain. (A) Representative image of whole embryo H&E at birth (P0) for *Bicra*^-/-^ pup and control heterozygous littermates (B) Representative image showing erythroblastic island (EBI) in fetal liver at birth for *Bicra*^-/-^ knockout pup and control *Bicra*^+/-^ littermates (C) Representative image showing aortic RBCs at birth for *Bicra*^-/-^ pup and control *Bicra*^+/-^ littermates (D) Quantification of red area in liver using ImageJ software. n = 2 independent biological replicate experiments with 5 representative images/replicate taken for quantification, statistical test performed using Prism software Welch test (E) Quantification of RBCs in major vessels of heart using ImageJ software. n = 2 independent biological replicate experiments with 5 representative images/replicate taken for quantification, statistical test performed using Prism software Welch test (F) Frequency of fetal liver erythrocytes (Ter119^+^) in mice of all 3 genotypes, Statistical testing is performed using Prism software with Ordinary 1-way ANOVA (with multiple comparison) (G) Representative flow plots for different stages of erythrocytes maturation using Ter119^+^ and CD71^+^ gated cells isolated from fetal liver at E14.5 (H) Quantification of different stages of erythrocytes maturation for *Bicra*^-/-^ pup and littermate control pups; statistical test performed using Ordinary 1-way ANOVA (with multiple comparison). A designation of * = P ≤ 0.05; ** = P ≤ 0.01; *** = P ≤ 0.001; ns = non-significant (P > 0.05).

To test whether the red blood cell defect arises from improper regulation of the *ß* globin gene locus, as was observed with *Smarca4* deletion (Bultman, Gebuhr, and Magnuson 2005), we performed RT-qPCR for different *ß* globin variants (*embryonic: Ey*, *ßh1; adult: ß major, ß minor*), but observed no differences in the expression of any *ß* globin variants across the genotypes (**Figure S2D**). The lineage marker Ter119 is expressed on early proerythroblasts all the way to mature erythrocytes in circulation; therefore, we first evaluated the number of erythroid cells expressing the lineage marker Ter119 in fetal livers. Using flow cytometry, we observed no statistically significant differences in the number of Ter119^+^ cells between different genotypes (**Figure 2F**). In addition, there was no significant change in the sub-populations representing the different stages of maturation of erythroid progenitors, as quantified by Ter119 and CD71 marker status (Koulnis et al. 2011) (**Figure 2G and 2H**). Colony forming assays with fetal liver cells from E14.5 pups showed no difference in CFU-E or CFU-GEMM between genotypes (**Figure S2E**), suggesting that early erythroid progenitors were not affected by *Bicra* loss. We did, however, observe a small but statistically significant decrease in the number of colony-forming units (CFUs) for CFU-GM (**Figure S2E**).

### Fetal livers from Bicra^-/-^ mice have reduced macrophage numbers and altered macrophage-specific gene expression

Since there was no observable defect in red blood cell-intrinsic factors or the expression of *ß* globin genes, we performed bulk RNA-Seq on E14.5 fetal livers to identify genes associated with *Bicra* deletion. We identified 108 differentially expressed genes (DEG) with >|1.5| fold change (*P*_adj_ < 0.05) in the *Bicra^-/-^*mice relative to *Bicra^+/+^* littermates (**Figure 3B**), where the majority of DEGs (97/108) were downregulated. The *Bicra^-/-^* samples clustered separately from both *Bicra^+/-^*and *Bicra^+/+^* samples, in agreement with the relatively normal phenotype for the *Bicra^+/-^* pups (**Figure 3A**). Similarly, gene expression changes for the *Bicra^-/-^* livers compared to either *Bicra^+/-^* or *Bicra^+/+^* livers were similar (**Figure S3A**). Nevertheless, *Bicra^+/-^* livers did show differential expression for a handful of genes (**Figure 3C, 3D and S3B**). Gene set enrichment analysis (GSEA) with MSigDB hallmark gene sets indicated a significant downregulation of inflammatory response and interferon response genes in the *Bicra^-/-^*livers (**Figure S3C**), and pathway analysis indicated that the downregulated genes in *Bicra^-/-^* livers are associated with acinar cells and macrophages (**Figure 3E**). Macrophage-specific genes (*Cd5l, Cybb, Csf2rb, Siglec1, Ccr2*) and lytic enzymes (*Pnliprp1, Pnliprp2, Cpb1, Cel, Prss2, Rnase1, Pnlip*) were downregulated in both *Bicra^-/-^*and *Bicra^+/-^* livers (**Figure 3F, Figure S3D**). Lytic enzymes are highly expressed in acinar cells, but also play a role in macrophage function (Baggiolini and Schnyder 1982). These macrophage-specific transcriptional changes prompted us to investigate whether the *Bicra^-/-^*livers have a reduced number of macrophages. Using F4/80 and CD11b as macrophage-specific markers, we performed flow cytometry to calculate the percentage of macrophages in the fetal liver for the genotypes (**Figure 3I, 3J**). We observed a significant reduction in the number of macrophages in homozygous knockout mice compared to both wildtype and heterozygous littermates. Further, IHC for F4/80 marker on fixed sections confirmed this reduction of macrophages in *Bicra^-/-^* livers (**Figure 3G, 3H**). Tissue-resident macrophages in an E14.5 stage embryo also reside in other organs, such as the brain, heart and lung (Hoeffel et al. 2015); however, the macrophage levels at these sites were not significantly affected by *Bicra* deletion (**Figure S3E, S3F, S3G, S3H**).

**Figure 3.**
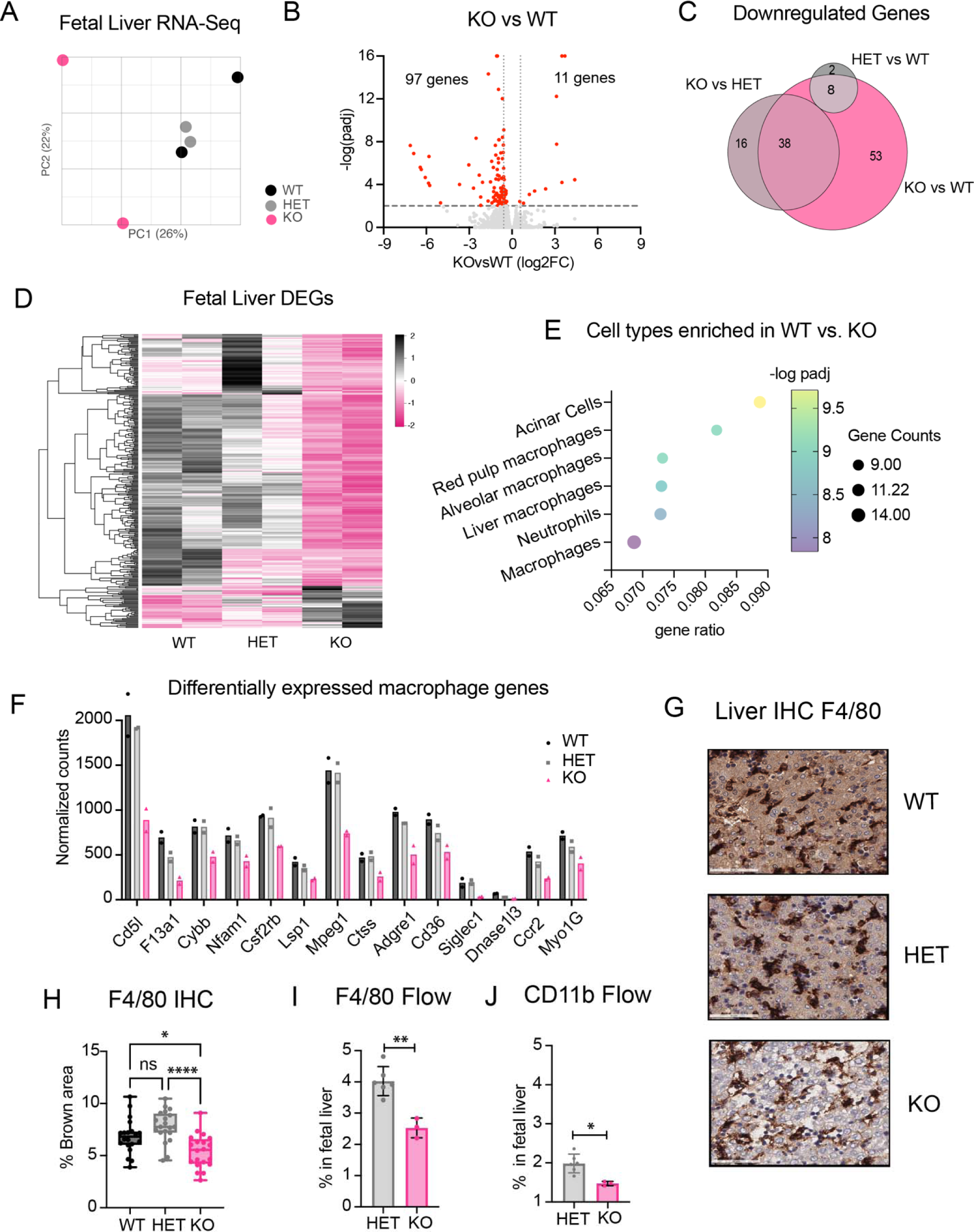
Fetal liver transcriptional analysis and macrophage frequency at embryonic stage 14.5. (A) Principal component analysis of bulk RNA-Seq samples isolated from fetal liver at E14.5. (B) Volcano plot of fetal liver RNA-Seq gene expression changes post *Bicra* homozygous knockout compared to wildtype littermate controls at E14.5. The log2 fold change indicates the mean expression level for each gene. Each dot represents one gene. The differentially expressed genes (DEGs) are shown in red with *p*_adj_ < 0.05 and FC > |1.5|. (C) Overlap of downregulated genes (*p*_adj_ < 0.05, FC > |1.5|) in *Bicra^-/-^* knockout mice compared to *Bicra^-+/+^* wildtype (bright pink), *Bicra^-/-^* knockout mice compared to *Bicra^-/+^* heterozygous mice (light pink) and *Bicra^-/+^* heterozygous mice compared to *Bicra^-+/+^* wildtype wildtype (grey). (D) Heatmap of differentially expressed genes identified as downregulated with *Bicra^-/-^* knockout using DESeq2 method. Normalized counts were calculated between different genotypes with two biological replicates. (E) The top six most significantly enriched cell types represented in the genes significantly decreased upon *Bicra* knockout. (F) Macrophage specific gene expression quantified using normalized counts between mice of different genotypes. (G) Representative image of fetal liver macrophage stained using F4/80 marker at E14.5 for *Bicra*^-/-^ pup, control littermates (H) Quantification of F4/80^+^ staining in E14.5 fetal liver using Image J software, n = 2 independent biological replicate experiments with 5 representative images/replicate taken for quantification, statistical test performed using Prism software using Ordinary 1-way ANOVA (with multiple comparison) (I) Frequency of F4/80^+^ macrophage using cell sorting, statistical test performed using Prism software Welch test (J) Frequency of CD11b^+^ macrophage using cell sorting, statistical test performed using Prism software Welch test. A designation of * = P ≤ 0.05; ** = P ≤ 0.01, *** = P ≤ 0.001, ns = non-significant (P > 0.05).

### Bicra^-/-^ mice have a defect in liver-resident macrophages

The phenotypic analysis pointed to a defect in red blood cells and the bulk RNA-Seq pointed to a reduction in macrophage-specific gene expression in *Bicra^-/-^* mice livers. Therefore, we next performed RNA-Seq on FACS-sorted F4/80^+^ macrophages and Ter119^+^ sorted erythrocytes to identify the genes specifically regulated by *Bicra* in these cell populations. In F4/80^+^ macrophages we identified 899 genes with decreased expression and 1203 genes with increased expression in the *Bicra^-/-^* livers compared to wildtype littermates (**Figure 4A)** (FDR < 0.05, fold change (FC) > |1.5|). In contrast, only three genes were differentially regulated in the Ter119^+^ erythrocytes enriched from knockout livers (**Figure S4A**), supporting a more substantial role for *Bicra* in gene regulation in the macrophages than erythrocytes. Similar to the bulk RNA-Seq, more genes were differentially expressed in *Bicra^-/-^*knockout macrophages relative to *Bicra^+/+^* wild type macrophages than were differentially regulated in *Bicra^+/-^* heterozygous macrophages relative to *Bicra^+/+^* wild type macrophages (**Figure S4B)**. Also similar to the bulk-RNA-Seq, the *Bicra^-/-^* samples clustered separately from *Bicra^+/-^* and *Bicra^+/+^*samples **(Figure S4C)** and gene expression changes for the *Bicra^-/-^* macrophages compared to either *Bicra^+/-^* or *Bicra^+/+^* macrophages were similar **(Figure S4D)**. Gene set enrichment analysis (GSEA) performed using cell-type specific gene sets showed strong positive enrichment of erythroid genes and negative enrichment of macrophage genes in *Bicra^-/-^* liver macrophages compared to *Bicra^+/+^*(**Figure 4B**). Supporting this, GSEA performed using Gene Ontology (GO) gene sets showed strong positive enrichment for erythrocyte development genes and negative enrichment for phagocytosis genes in the *Bicra^-/-^* liver macrophages compared to *Bicra^+/+^* WT (**Figure S4E**) and GO pathways enriched in the genes upregulated in the *Bicra^-/-^* macrophages are associated with extracellular matrix organization and processes involving cell-cell adhesion, while the genes downregulated in *Bicra*^-/-^ macrophages are enriched for pathways related to inflammation and immune activation (**Figure 4C)**.

**Figure 4.**
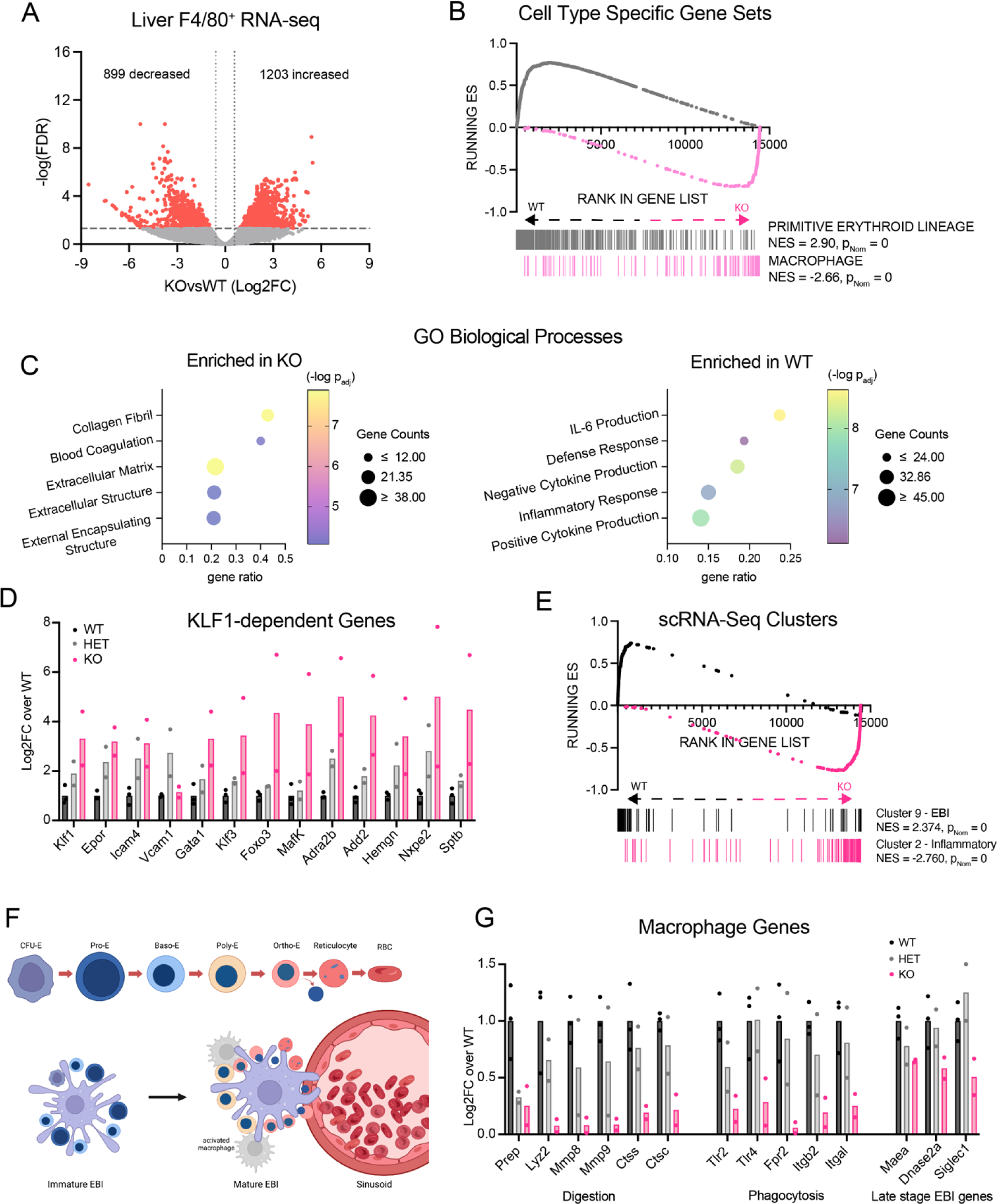
(A) Volcano plot of fetal liver sorted F4/80^+^ macrophage RNA-Seq gene expression changes post *Bicra* homozygous knockout compared to wildtype littermate controls at E14.5. The Log2 fold change indicates the mean expression level for each gene. Each dot represents one gene. The differentially expressed genes are shown in red with FDR < 0.05 and FC > |1.5|. (B) GSEA of RNA-Seq from fetal liver F4/80^+^ sorted macrophages from wildtype and knockout mice. (C) The top five most significantly enriched cell-type represented in the genes significantly increased (left) and decreased (right) upon *Bicra* knockout. Analysis performed using Enrichr. (D) KLF1-dependent gene expression depicted as the fold change (FC) of the average of the normalized counts for *Bicra^-+/+^*wildtype samples. (E) GSEA of RNA-Seq from fetal liver F4/80^+^ sorted macrophages from wildtype and knockout mice. Gene sets were created from the top 100 unique genes from each cluster defined using scRNA-Seq data of mouse fetal (E14.5) liver macrophages (Kayvanjoo et al. 2023). (F) The immature and mature EBIs associated with early and late erythrocyte development, respectively (G) Expression of genes associated with mature EBI function depicted as the fold change (FC) of the average of the normalized counts for *Bicra^-+/+^*wildtype samples.

The central macrophage in erythroblastic islands directly interacts with erythroblasts throughout all stages of maturation and at the end, engulfs the nuclei and releases mature red blood cells into blood vessels (Palis 2017; Kayvanjoo et al. 2024). At early stages, EBI macrophages adhere tightly to erythroblasts, and will co-enrich erythroblasts or portions of erythroblasts during FACS (Popescu et al. 2019). In addition, the EBI macrophages themselves express a subset of erythrocyte-specific genes (Kayvanjoo et al. 2024; Mukherjee et al. 2021). In our RNA-Seq data, *Bicra*^-/-^ macrophages are significantly enriched (>4 fold) for erythrocyte-specific genes, such as *Hb* hemoglobin genes, *Sptb*, *Trim10*, *Nxpe2*, *Snca*, and *Epb42* (**Figure S4F**). *Bicra*^-/-^ macrophages also have increased expression of *Klf1*, the defining transcription factor for EBI macrophages (Porcu et al. 2011) as well as KLF1-dependent genes in fetal liver macrophages, such as *Epor, Icam4*, *Adra2b*, and *Add2* (Mukherjee et al. 2021) (**Figure 4D**). This indicates that in the *Bicra*^-/-^ liver, genes associated with EBIs are actually enriched in the macrophage population.

While EBI macrophages are considered the most critical macrophage for erythrocyte maturation, they are part of a heterogeneous population of macrophages in the fetal liver (Mukherjee et al. 2021; Seu et al. 2017). To further identify which specific macrophage populations are affected in *Bicra^-/-^*, we used a recently published scRNA-Seq dataset from E14.5 mouse livers (Kayvanjoo et al. 2024). They identified 6 clusters with macrophage-specific gene expression and characterized the unique gene expression differences between the clusters. Using the top 100 genes identified in each cluster (Kayvanjoo et al. 2024), we performed GSEA with our RNA-Seq data from F4/80 sorted macrophages. Genes from macrophage cluster 9, which was defined as EBI macrophages, were positively enriched in *Bicra*^-/-^ macrophages, while genes from macrophage cluster 2, which were defined as inflammatory macrophages, were the most negatively enriched in *Bicra*^-/-^ macrophages (**Figure 4E**). Clusters 1 and 11 that represent macrophage and granulocytic precursors, respectively, were also negatively enriched in *Bicra*^-/-^ macrophages (**Figure S4G**). This supports the hypothesis that *Bicra*^-/-^ livers have an increased percentage of EBI macrophages, but a significantly decreased percentage of inflammatory macrophages. Supporting this, there is a notable decrease (>10x) in the expression of genes associated with inflammatory macrophages, such as *Itgam*, *Csf1r*, *Il4ra*, *Ccr2*, *Mpeg1*, *Ly6c2*, and *S100a8/9* (**Figure S4H**). Since ∼30% of liver macrophages fall into the scRNA-Seq inflammatory gene cluster 2 (Kayvanjoo et al. 2024), the depletion of this population could account for the overall decrease in F4/80^+^ cells observed in the *Bicra*^-/-^ livers (**Figure 3I**). While there has not been a reported role for inflammatory macrophages in erythrocyte maturation, inflammatory Cd11b^+^ (*Itgam*) macrophages are present in erythroblastic islands along with the central erythrocyte-bound macrophage (Seu et al. 2017). Since our RNA-Seq in *Bicra^-/-^* macrophages indicates that immature EBI macrophages are present and able to bind erythrocytes, we next evaluated whether there is a defect in genes specific to the mature EBI macrophages that are involved in enucleation, phagocytosis, and digestion during the final RBC maturation steps (**Figure 4F**). In the *Bicra^-/-^* liver macrophages, genes involved in protein digestion and phagocytosis are decreased (**Figure 4G)**, although there is no way to distinguish whether this gene expression decrease is merely a reflection of the decrease in inflammatory macrophages. Therefore, we looked at genes specifically expressed by, and required for, mature phagocytotic EBI macrophages, such *Maea* (Emp) (Soni et al. 2006), *Dnase2a* (Kawane et al. 2001) and *Siglec1* (Cd169) (Chow et al. 2013) (**Figure 4G**). All three genes are significantly decreased in the *Bicra^-/-^*liver macrophages, indicating a possible defect in mature EBI macrophage function. Emp (*Maea*) is an adhesion protein expressed on both EBI macrophages and erythrocytes that is required for nuclear extrusion and enucleation of erythrocytes (Soni et al. 2006). The preweaning lethality and the red blood cell defect phenotype we observe for the *Bicra*^-/-^ mouse most closely mirrors the reported phenotypes observed in the Emp knockout mouse (Soni et al. 2006); therefore, we next evaluated late-stage erythrocyte maturation and enucleation in the *Bicra^-/-^* knockout mice.

### Red blood cell maturation is defective in *Bicra^-/-^* mice

To evaluate whether late-stage erythrocyte maturation and enucleation are defective in knockout mice, we evaluated Wright-Giemsa-stained fetal liver cytospins from E14.5 embryos. *Bicra^+/+^* wildtype fetal livers had approximately 10-20% nucleated erythrocytes, which is similar to what has been observed previously (Soni et al. 2006). The *Bicra^-/-^* knockout mice had higher proportions of nucleated erythroblasts (30-50%) with fewer enucleated erythrocytes (**Figure 5A).** In agreement with these results, the number of immature nucleated erythrocytes were increased in blood smears from E18.5 *Bicra^-/-^* embryos (**Figure 5B**) and in paraffin-embedded fetal liver sections of E18.5 pups (**Figure S5A**). Since the heterozygotes had a similar, although much more mild transcriptional profile compared to homozygotes, we performed a blood profile from 6-month-old adult *Bicra^-/+^* heterozygous and *Bicra^+/+^* wild type mice to identify if there were any defects in adult erythrocyte maturation. The only difference was a slight, but not statistically significant (p = 0.09), increase in the mean corpuscular volume in the heterozygotes (**Figure S5B**). This is consistent with the phenotype reported for *Bicra*^+/-^ mice by KOMP (Dickinson et al. 2016; Groza et al. 2023) and could indicate a mild defect in RBC maturation that leads to an increase in the slightly larger immature reticulocytes (J. Liu et al. 2010). All in all, the data supports a role for *Bicra* in red blood cell maturation and enucleation by fetal liver macrophages.

**Figure 5.**
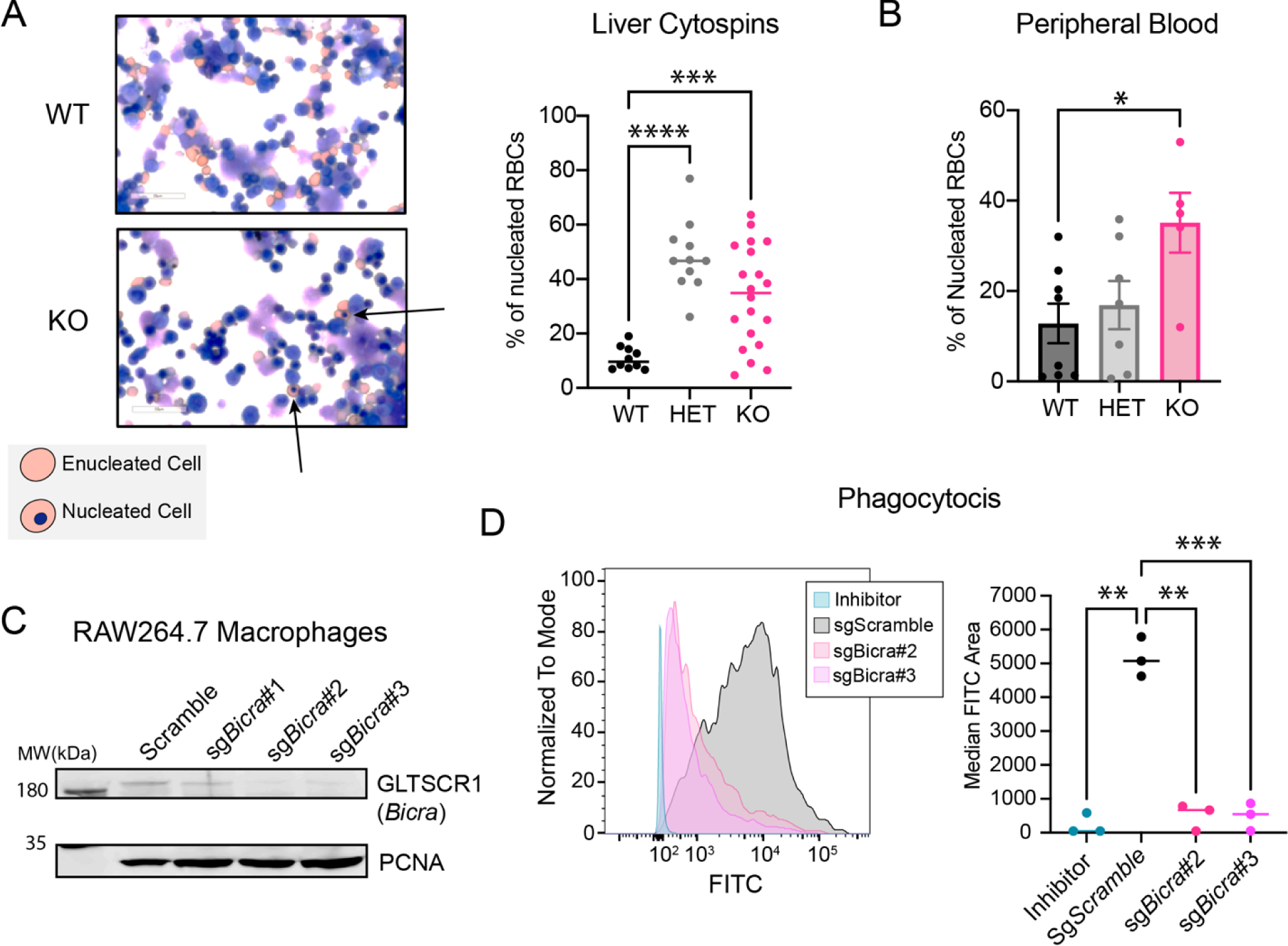
(A) Representative image of cytospin preparation from fetal liver at E14.5 for *Bicra*^-/-^ knockout pup and control homozygous littermates (left) Quantification of nucleated red blood cells in fetal liver cytospin preparation at E14.5 for *Bicra*^-/-^ knockout pup and control littermates using Image J software, n = 3 independent biological replicate experiments with 5 representative images/replicate taken for quantification, statistical test performed using Prism software using Ordinary 1-way ANOVA (with multiple comparison) (right). (B) Quantification of nucleated red blood cells in peripheral blood at E18.5 for *Bicra*^-/-^ knockout pup and control littermates using Image J software, n >/=6 independent biological replicate experiments with 5 representative images/replicate taken for quantification, statistical test performed using Prism software using Ordinary 1-way ANOVA (with multiple comparison). (C) Protein lysates isolated from RAW cell line; PCNA is used as a loading control. (D) Quantification of median FITC area for the scramble control, inhibitor treated and knockout cell line; Statistical testing is performed using Prism software with using Ordinary 1-way ANOVA (with multiple comparison). (E) Flow cytometry histogram comparing the scramble control (grey), inhibitor (cyan), sgBicra#2 (bright red-pink), sgBicra#3 (fuchia). A designation of * = P ≤ 0.05; ** = P ≤ 0.01; *** = P ≤ 0.001; ns = non-significant (P > 0.05).

### Bicra^-/-^ knockout in RAW 264.7 cells display reduced phagocytosis

To evaluate if the phagocytic function of macrophages is disrupted by the loss of *Bicra,* we performed a phagocytosis assay using the mouse macrophage cell line RAW 264.7 (Raschke et al. 1978) and FITC-conjugated IgG beads. We generated *Bicra* knockout cell lines using two separate guide RNAs that display high (>90%) knockout efficiency (**Figure 5C**). We observed a statistically significant reduction in the phagocytic ability of *Bicra*^-/-^ cells compared to scramble control (**Figure 5D, Supplemental video 2** and **Supplemental video 3**), which was similar to that of cytochalasin D, a chemical inhibitor of phagocytosis (**Supplemental video 4**) (Magae et al. 1994). Comparative analysis of flow cytometry histograms between the scramble control and each knockout cell line/inhibitor showed a left shift indicative of reduced phagocytic activity (**Figure 5D**). This *in vitro* assay may explain the accumulation of nucleated RBCs due to a reduction in the phagocytic capability of macrophages in *Bicra*^-/-^ mice. Potentially related to the reduction of inflammatory macrophages *in vivo* in the livers of *Bicra*^-/-^ knockout mouse, loss of *Bicra* in the phagocytic macrophage cell line (RAW 264.7) reduced overall cell number (**Figure S5D**).

## Discussion

Hematopoiesis is a stepwise lineage commitment process that generates the numerous mature blood cell types (Orkin and Zon 2008). The undifferentiated multipotent hematopoietic stem cells progressively mature into lymphoid, myeloid, and erythroid lineages (Weissman 2000). This hierarchical commitment involves dynamic crosstalk between lineage specific transcription factors and epigenetic regulators such as the BAF complex (Graf and Enver 2009; Ho and Crabtree 2010). Many subunits of the BAF complex are required for various aspects of hematopoiesis (Z. Tu and Zheng 2022), including shared subunits SMARCA4 (J. Tu et al. 2020), SMARCA2 (Naidu et al. 2022), SMARCC1(Wu et al. 2020), and ACTL6A (Krasteva et al. 2012), as well as cBAF specific subunits ARID1A (L. Han et al. 2019; Krosl et al. 2010), ARID1B (Madan et al. 2023) and DPF2 (Mas et al. 2023), PBAF-specific subunits ARID2 (L. Liu et al. 2018; Bluemn et al. 2021) and PHF10 (Krasteva, Crabtree, and Lessard 2017), and GBAF-specific subunit BRD9 (Xiao et al. 2023; Garg et al. 2024). The contribution of the remaining subunits is yet to be explored.

BICRA/GLTSCR1 is a unique complex-defining subunit of GBAF/ncBAF complex in both human and mouse cell lines (Alpsoy and Dykhuizen 2018; Sandoval et al. 2018; Gatchalian et al. 2018); however, many of its functions are redundant with its mutually exclusive paralog, BICRAL/GLTSCR1L. Its unique biological function at an organismal level remains unclear. The paralog-specific roles for GLTSCR1 are likely related to GBAF function in general, which has been explored via targeting the bromodomain of BRD9, another unique subunit required for GLTSCR1 incorporation and stability (Alpsoy et al. 2021). BRD9 is required for embryonic stem cell gene expression (Gatchalian et al. 2018), preventing myelodysplastic syndrome (MDS) (Inoue et al. 2019), promoting leukemia (Del Gaudio et al. 2019), and macrophage gene activation (L. Wang et al. 2021; Ahmed et al. 2022). Specifically in hematopoiesis, conditional deletion of *Brd9* in HSCs results in a bias towards myeloid cell lineage (monocytes and neutrophils but not erythrocytes) both in human cord blood cells and in mice *in vivo* (Xiao et al. 2023). Reduction of BRD9 in human HSPCs is similar, with enhanced terminal myeloid differentiation but reduced megakaryocytic and erythroid differentiation (Garg et al. 2024; Lara-Astiaso et al. 2023). These indicate a general role for GBAF in myeloid cell development and function, but the relative contributions of GLTSCR1 and its paralog GLTSCR1L are unclear. To address the *in vivo* function of GLTSCR1 we generated a mouse lacking functional endogenous *Bicra* using CRISPR-Cas9. Heterozygous mice were born healthy and fertile, whereas all the homozygous mice died within 24 hours after birth with anemia. Histology and flow cytometry analysis of the fetal liver identified a slight reduction in liver resident macrophages, while RNA-Seq revealed aberrant macrophage gene expression. In addition, we identified a slight defect in progenitor potential in *Bicra* knockout mice, with a decrease in CFU-GM colonies that produce myeloid cells. Consistent with our findings, an epigenetic CRISPR knockout screen identified reduced myeloid priming upon loss of *Bicra* in irradiated mice 14 days post transplantation with LSK multipotent (Lin−Sca1+c-Kit+) progenitor cells (Lara-Astiaso et al. 2023). Secondly, in line with our findings, loss of *Brd9* (another defining subunit of GBAF), also shows myeloid-lineage skewing and an increase in mean corpuscular volume (MCV) (Xiao et al. 2023), suggesting that GBAF regulates myeloid cell specification. This could be due to a role for GLTSCR1 in the progenitors themselves and reflected by the decrease in CFU-GM (**Figure S2E**). Alternatively, the decrease in CFU-GM could be due to a defect in the fetal liver macrophages. Deletion of all resident fetal liver macrophages results in a similar CFU-GM decrease (Kayvanjoo et al. 2024), In addition, fetal liver macrophages are required for granulopoiesis (Romano et al. 2022), genes of which are decreased in the *Bicra*^-/-^ liver macrophage RNA-Seq (**Figure S4F**). Conditional knockouts will help define the intrinsic vs macrophage-driven effects hematopoietic progenitors.

The perinatal lethal phenotype we observed is different than what was reported previously for *Bicra* knockout mice created using CRISPR-mediated exon 5 disruption (F. Han et al. 2023). In this study the authors reported mid to late gestation embryonic lethality due to a heart defect. However, neither the conditional cardiac nor neural crest-specific *Bicra* deletion resulted in embryonic lethality, complicating the interpretation that the heart defect is responsible for the embryonic lethality. One potential caveat of our *Bicra* knockout is that in some cell types we observe the expression of a small amount of a GLTSCR1 variant stemming from an alternative start site at exon 6 (Smith et al. 2000). This variant is expressed at low levels and is missing the conserved N-terminal domain but can still associate with the GBAF complex (**Figure S5E)**. It is unlikely that this variant can account for the difference in phenotypes as the *Bicra^-/-^* knockout mouse produced by the IMPC initiative (Dickinson et al. 2016; Groza et al. 2023) was generated using CRISPR-mediated deletion in exon 7-10 (Elrick et al. 2021) and also exhibits preweaning lethality for *Bicra* homozygous knockout pups and increased mean corpuscular volume (MCV) in *Bicra* heterozygous pups. In agreement with the previous publication, they also noted a heart hemorrhage, specifically in the *Bicra* knockout female pups. Our H&E sections don’t indicate any gross morphological difference in the heart of *Bicra* knockout mice; however, they aren’t of sufficient detail to detect the reported wall thinning (**Figure S5F**). While the heart defect can’t be ruled out as a contributing factor to the mortality of *Bicra* knockout mice, the macrophage defect we observe is consistent with the pale appearance and preweaning lethality.

Our results indicate that *Bicra* is essential for liver macrophage function during development, and that its paralog *Bicral* is unable to fully compensate; however, it doesn’t address whether *Bicral* is similarly required in liver macrophages. There is no published manuscript on a *Bicral* knockout mice but the mouse generated through the International Mouse Phenotyping Consortium (Dickinson et al. 2016) using CRISPR-mediated deletion of exon 2 and 3 is viable with bone defects; however, the red blood cells do have increased corpuscular volume, which could indicate some contribution of *Bicral* in macrophage function during erythrocyte maturation.

Previous studies defining the importance of EBI macrophages during development have focused primarily on the adhesion molecules that establish contacts between the EBI macrophage and erythrocytes, most notably, Erythroblast Macrophage Protein (EMP) (Hanspal and Hanspal 1994), VCAM1 (Sadahira, Yoshino, and Monobe 1995), and ICAM-4 (Lee et al. 2006) which are all required for erythrocyte maturation and enucleation. Perhaps the most well-studied protein in erythrocyte maturation is EMP, encoded by *Maea*. Both EBI macrophages and erythrocytes express EMP and their crosstalk with developing erythroblasts helps in the enucleation and maturation of functional red blood cells (Skutelsky and Danon 1972; Hanspal and Hanspal 1994; Hanspal, Smockova, and Uong 1998). *Maea*^-/-^ homozygous knockout mice die soon after birth with anemia owing to the reduced number of terminally mature red blood cells (Soni et al. 2006) and the resulting phenotypes of germline *Maea*^-/-^ knockout mice most closely resemble those with germline *Bicra*^-/-^ knockout mice. Other factors required for erythrocyte maturation include *Dnase2a*, which is regulated by KLF1 in EBI macrophages (Porcu et al. 2011) and is required for nuclei digestion after phagocytosis. Our F4/80^+^ sorted-macrophage RNA-Seq identified an increase in genes associated with EBI macrophages and their associated erythrocyte cells. Many genes specifically required for EBI macrophage-erythrocyte adhesion are increased or unaffected in the *Bicra^-/-^*macrophages, such as *Klf1*, *Icam4*, *Vcam1*, *Mertk*, *Spic*, *Igf1, Scl40a1*, *Trf*, *Timd4*, *Vegfb*, *Il33*, *Sptb*, and *Add2.* This indicates that adhesion to erythrocytes is not impacted by *Bicra*/GLTSCR1. EBI macrophages from mice with knockout of EMP (*Maea*) similarly adhered normally to erythrocytes but were unable to enucleate them (Soni et al. 2006). Similarly, genes required for the enucleation process were decreased in the *Bicra^-/-^* macrophages, including *Itgav*, *Maea*, *Dnase2*a, *Siglec1*, *Adgre1*, *Il18*, *Il4ra* (Li et al. 2021; Mukherjee and Bieker 2021). In addition, *Ly6c*, which is expressed only on a subset of EBI macrophages (Li et al. 2019) was decreased, although its role in enucleation isn’t known. This indicates that there is either a reduction in the subset of EBI macrophages required for enucleation, or a block in the ability of EBI macrophages to be activated to a phagocytic state for enucleation.

The most prominent effect of *Bicra^-/-^* knockout on fetal liver macrophages is the reduction in genes associated with inflammatory macrophages, which are separate from EBIs. Imaging studies indicate that inflammatory macrophages are not physically associated with erythrocytes (Kayvanjoo et al. 2023); however, inflammatory CD11b+ (*Itgam*) macrophages, are present in the EBI niche (Seu et al. 2017). While these inflammatory macrophages are considered unimportant for <u>formation</u> of EBIs (Mukherjee and Bieker 2021); their role in EBI <u>function</u> has not been tested. The most robustly decreased genes in the *Bicra^-/-^* macrophages are Spi1/PU.1 target genes. PU.1 is known to associate with SWI/SNF complexes (Chambers et al. 2023; Frederick et al. 2023) and be required for monocyte development and innate immunity (Zhao et al. 2022; Iwasaki et al. 2005). In addition, *Spi1* knockout mice have lower numbers of enucleated RBCs (Kayvanjoo et al. 2023), although they also have 90% reduction of all macrophages in the fetal liver, which complicates the interpretation somewhat. Our data suggests that the inflammatory macrophages could be required for providing instructional signals to EBI macrophages that facilitate the last steps of erythrocyte enucleation and maturation. Alternatively, *Bicra* is required for inflammatory gene expression in both macrophage subsets. Additional experiments will be required to elucidate the relevant macrophage type and function dependent on *Bicra*/GLTSCR1. In subsequent studies it will be beneficial to employ inducible Cre-loxP systems to evaluate if *Bicra* deletion in the macrophage compartment alone fully recapitulates the perinatal lethality and erythrocyte defect, and then to further define which macrophage subsets require *Bicra* for differentiation and function during development.

## Material and methods

### Generation of Bicra knockout mice

The transgenic mice were generated in collaboration with the Purdue University Transgenic Animal Core Facility. Fertilized mouse embryos were harvested from C57BL/6J (CD45.2^+^) mice at the blastocyst stage and co-injected with RNA encoding *Cas9* and sgRNA (CCCCATTGTCAGCGGGCTGT) construct targeting exon 4 of *Bicra*. The embryos were implanted in pseudopregnant C57BL/6J females. After delivery and weaning, the pups were genotyped for the presence of the edited allele at the CRISPR target site using semi-quantitative PCR and Bicra deletion was further confirmed through Sanger Sequencing. For the F_0_ offspring, a *Bicra*^+/-^ heterozygous male with 8bp deletion in a single allele was backcrossed five times to wildtype C57BL/6J (CD45.2^+^) females. The *Bicra*^+/-^ male mouse was also backcrossed with 129SvE female.

### Animal husbandry and genotyping

Animals were housed in a specific pathogen free animal facility and handled in accordance with the guidelines of Purdue Institutional Animal Care and Use Committee (IACUC). All experimental procedures were approved by the IACUC. Tail clips were taken from each pup on the weaning day and genotyping was performed using the GoTaq® Flexi DNA Polymerase kit (Promega). Separate PCR reactions were set up to amplify either the wildtype or mutant allele using forward wildtype primer (5’-CCCATTGTCAGCGGGCTGTAG-3’), forward mutant primer (5’-GCCCCATTGTCAGCGGGTG-3’), and universal reverse primer (5’-CAGCGTGGACCTGGACTTCCTAG-3’).

### Protein isolation from mice liver tissue

Pregnant mice were euthanized using isoflurane asphyxiation followed by cervical dislocation as recommended by the American Veterinary Medical Association’s (AVMA) guidelines 2020. *In utero* mouse pups were isolated and each pup’s liver tissue was obtained and homogenized in an ice-cold buffer A (20 mM HEPES, pH 7.9, 25 mM KCl, 10% glycerol with PMSF, aprotinin, leupeptin, and pepstatin) using a 27-gauge needle. The nuclei were pelleted by centrifugation at 600 × *g* (Eppendorf Centrifuge 5810 R, Hamburg, Germany) for 5 minutes. The nuclei pellet was resuspended in Buffer C (20 mM HEPES, pH 7.9, 100 mM KCl, 3 mM MgCl_2_, 0.1 mM EDTA, 10% glycerol with PMSF, aprotinin, leupeptin, pepstatin and benzonase) and incubated at room temperature for 20 minutes. Sequential protein precipitation was performed by adding 3 M ammonium sulfate to the suspension followed by incubation in the cold room for 30 minutes. The proteins were sub-fractionated using ultracentrifugation at 100,000 × *g* for 30 minutes. The supernatant was then transferred to a new tube and 0.3 g/mL of ammonium sulfate powder was added for salting out the proteins followed by centrifugation at 21,000 × *g* for 30 minutes. The protein pellet was redissolved in chromatin buffer (20 mM HEPES, pH 7.9, 150 mM NaCl, 1%Triton X-100, 7.5 mM MgCl_2_, 0.1 mM CaCl_2_).

### Western blot analysis

Isolated proteins from nuclear extracts either from cell pellet or whole tissue were quantified using Pierce^TM^ Protein Assay Kit (Thermo Scientific, Rockford, IL). Each sample protein was supplemented with 4× lithium dodecyl sulfate (LDS) sample buffer containing 10% 2-mercaptoethanol followed by denaturation at 95 °C for 5 minutes. The proteins were separated on a 4–12% SDS-polyacrylamide gel and transferred to a PVDF membrane (Immobilon FL, EMD Millipore, Billerica, MA). The membrane was blocked using Immobilon (Immobilon® Signal Enhancer, EMD Millipore Corp., Burlington, MA) for 1 hour and incubated with primary antibodies overnight at 4 °C. The primary antibodies were detected by incubating the membranes with goat anti-rabbit or goat anti-mouse secondary antibodies (LI-COR Biotechnology, Lincoln, NE) conjugated to IRDye 800CW or IRDye 680, respectively, at room temperature for 1 hour. The IR signals were visualized using an Odyssey Clx Imager (LI-COR Biotechnology).

### Immunoprecipitation

Cells were harvested by trypsinization (0.25% Trypsin, Corning Mediatech, Inc.) and washed twice in ice-cold phosphate buffered saline (pH 7.2) followed by centrifugation at 250 × *g* for 5 minutes. The cell pellet was resuspended in buffer A (20 mM HEPES, pH 7.9, 25 mM KCl, 10% glycerol, 0.1% Nonidet P-40 supplemented with PMSF, aprotinin, leupeptin, and pepstatin) at a concentration of 20 million cells/mL. The cells were incubated on ice for 5 minutes, and the nuclei were isolated by centrifugation at 600 × *g* (Eppendorf Centrifuge 5810 R, Hamburg, Germany) for 10 minutes. The nuclei pellet was resuspended in chromatin IP buffer (20 mM HEPES, pH 7.9, 150 mM NaCl, 1%Triton X-100, 7.5 mM MgCl_2_, 0.1 mM CaCl_2_ supplemented with PMSF, aprotinin, leupeptin, and pepstatin). Additionally, 4 units/mL Turbo DNase (Ambion, Inc., Foster City, CA), was added to the extracts and rotated at 4 °C for 30 minutes. The protein extracts were spun down using centrifugation (Centrifuge 5424 R; Eppendorf, Hamburg, Germany) at 21,000 × *g* for 30 minutes. Two micrograms respective IgG was used per 0.3 mg lysate for immunoprecipitation. The immunocomplexes were captured using protein A/G magnetic beads (Dynabeads™ from Invitrogen™) following a two-hour incubation. The beads were washed twice in chromatin IP buffer and three times in high stringency wash buffer (20 mM HEPES, pH 7.9, 500 mM NaCl, 1% Triton X-100, 0.5% sodium deoxycholate, 1 mM EDTA). The proteins were eluted in 1× lithium dodecyl sulfate loading dye (ThermoScientific) by boiling at 70 °C for 10 minutes and then separated on a gel.

### Cell line maintenance and generation of CRISPR/Cas9-mediated knockout

RAW 264.7 parental cells were acquired from ATCC and were maintained in DMEM (Corning Mediatech, Inc.) supplemented with 10% heat-inactivated fetal bovine serum (Corning Mediatech, Inc.), 2 mM L-alanyl-L-glutamine (Corning Glutagro^TM^; Corning Mediatech), 1 mM sodium pyruvate (Corning Mediatech) and 100 units/mL penicillin and 100 g/mL streptomycin (Corning Mediatech). Cells were cultured in humidified incubator maintained at 37°C and 5% CO_2_. Cells were regularly tested for *Mycoplasma* contamination using MycoAlert Mycoplasma Detection kit (Lonza). Short guide RNA sequences targeting the mouse *Bicra* gene were retrieved and cloned into plenticrispr v2.0 (a gift from Feng Zhang Addgene plasmid no. 52961), and the vector was packaged into lentivirus using HEK293T cells as described in (Alpsoy and Dykhuizen 2018). RAW 264.7 cells were transduced with concentrated virus, and stable knockout lines were generated by puromycin selection.

### Histological analysis and Immunochemistry

Four C57BL/6 strain mice pups [two homozygous *Bicra^-/-^* mice and two heterozygous *Bicra^+/-^*littermate controls] and six 129SvE strain mice pups [two homozygous *Bicra^-/-^* mice, two heterozygous *Bicra^+/-^*, and two wildtype *Bicra^+/+^* littermate controls] were euthanized at birth via decapitation as recommended by AVMA. The whole pups were fixed for 24 hours in 10% neutral buffered formalin and dehydrated in 70% ethanol. Sample embedding, sectioning and H&E staining were performed by AML Laboratories (St. Augustine) and Histowiz Inc. (Long Island City). Fixed sections were stained for F4/80. For blood smears, peripheral blood was collected from freshly dissected embryos and spread on a clean beveled edge slide. The blood smears were air dried for 5 minutes, fixed using methanol for 2 minutes, and stained with HEMA 3^®^ STAT PACK (Fisher Scientific Company L.L.C. Kalamazoo, MI). Freshly isolated livers from timed pregnancies were used for cytospin preparation and stained with Wright-Giemsa stain (FisherBrand^TM^). Stained sections were visualized by using a Leica Aperio system at 40× magnification and staining quantification was performed with ImageJ 13.0.6 software.

### Methyl cellulose assay

After setting timed breeding, fresh embryos were harvested at E14.5. The embryos were euthanized by decapitation as recommended by AVMA guidelines. After liver dissection, single cell suspension was prepared by passing the cells through 70 µm nylon strainer. After filtration, cells were washed in 15 mL centrifuge tubes with IMDM/2% FBS (Corning Mediatech, Inc.) by a gentle spin at 300 × *g* for 5 minutes. The supernatant was removed, and cells were gently resuspended in IMDM/2% FBS. Single cell suspension was counted for each embryo. Cells were then plated in duplicates at 3,000 viable cells/plate using MethoCult methylcellulose media (R&D Systems, HSC007). After 14 days, plates were scored for colony counts using a light microscope (Olympus IX73) at 20× magnification.

### Flow cytometry

Freshly isolated fetal liver cells (E14.5 stage) were manually dissociated to a mononuclear cell suspension. The cells were washed with cold staining buffer (1× PBS, 0.2% BSA and 5 mM glucose) and spun down at 150 × *g* with low brake (Eppendorf Centrifuge 5810 R, Hamburg, Germany). The cell number was counted using a hemocytometer, and 1 million cells were blocked with PURE MS CD16/CD32 (Tonbo Bioscience) for 20 minutes. In all experiments dead cells were stained by using Fixable viability dye eFlour^TM^ 780 (ThermoFisher Scientific; Cat 65-0865). Surface markers for immune cells of interest were stained for 30 minutes in the dark. Gating thresholds were set to eliminate non-viable cells and separate multiple events (populations S0-S4/5 in fetal liver cells). Overlapping fluorescence from multiple antibodies was compensated using AbC Total Compensation beads (Thermo Fisher Scientific, catalog no. A10497). All the antibodies used in the experiments are summarized in Table T1. Flow cytometry was performed on Fortessa (BD Biosciences), and flow sorting was performed on FACSAriaII (BD Biosciences). Data were analyzed by FlowJo software (Tree Star, Ashland, OR).

### Quantitative real time PCR

For targeted gene expression, RNA was isolated from fetal liver cells using TRIzol reagent (Life Technologies Corporation, Grand Island, NY). The RNA purity was estimated using NanoDrop^TM^ (Thermo Fisher Scientific) spectrophotometer and cDNA synthesis was performed using Verso cDNA Synthesis Kit (Thermo Fisher Scientific, Baltics, UAB). RT-qPCR analyses were completed with PowerUP^TM^ SYBR Green Master Mix (Thermo Fisher Scientific, Baltics, UAB) using a Bio-Rad CFX96 Connect Real-Time System (Biorad Laboratories, Inc.). The primer sequences have been detailed in Table 2. *Rpl13*a was used for normalization of gene expression levels. The relative gene expression was calculated by 2^−ΔΔCt^, where Ct is the threshold cycle value (Livak and Schmittgen 2001).

**Table 2:**
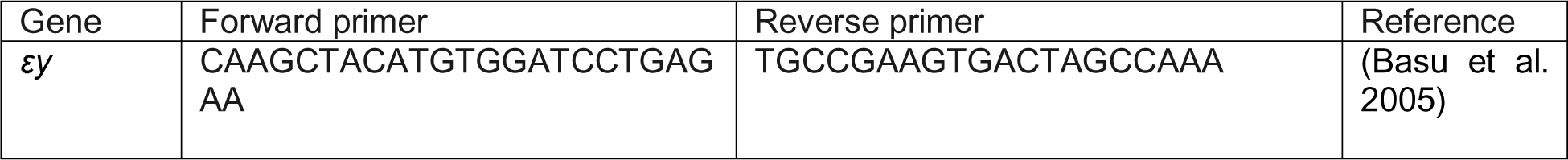
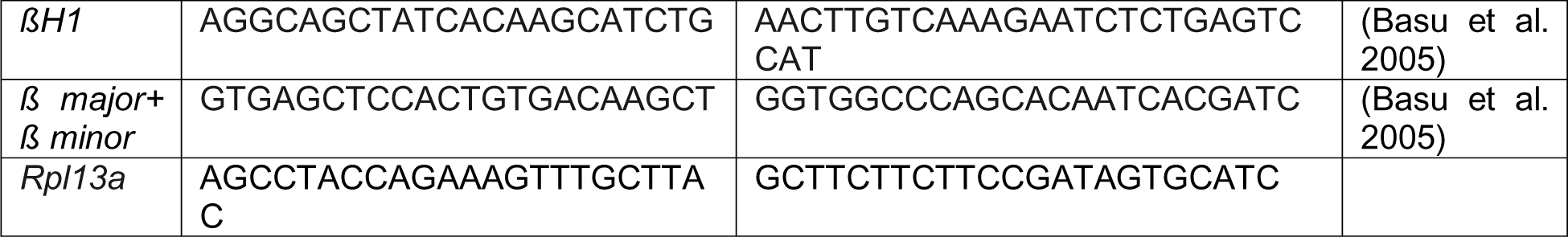

### Bulk RNA sample sequencing and data processing

Bulk RNA was isolated from freshly dissected fetal liver cells at E14.5 using TRIzol reagent (Life Technologies Corporation, Grand Island, NY). RNA samples were sequenced using Illumina sequencer following paired-end protocol and targeted read-length of 150 bp and >30 million total reads per sample. Data analysis was performed in collaboration with Purdue Computational Genomics Shared Resource. Quality trimming and filtering (adapter trimming, minimum Phred score 30, minimum read length 50 bp) was performed using Trim Galore v0.6.4. Quality trimmed reads were mapped against the mouse reference genome (mm10) using STAR aligner v2.5.4b. Number of reads aligned to genes were quantified with HTSeq package v0.7.0. Differential gene expression analysis was performed using DESeq2 v1.22.2. The Benjamini-Hochberg method was used to adjust the *P*-values for multiple testing. Genes with cutoff (*p*_adj_ < 0.05 and FC > |1.5|) were denoted as Differentially Expressed Genes (DEGs). Pathway analysis was performed using Enrichr (Chen et al. 2013). Enrichments of gene sets were performed using GSEA (Subramanian et al. 2005). Quantitative overlap analysis was performed using Eulerr (Larsson et al. 2024).

### Liver sorted macrophage and erythrocyte RNA Sequencing and data processing

Single cell suspension of fetal liver cells isolated from E14.5 embryos was prepared as mentioned in flow cytometry section. Cells were stained with anti-mouse F4/80 (Biolegend; Cat # 123122) and Ter119 antibodies. Stained cells were sorted on FACSAriaII (BD Biosciences) in PBS. Total RNA from cells was isolated using RNeasy Plus Micro Kit (Qiagen Inc., Valencia, CA) using the manufacturer protocol. RNA integrity and purity was assessed on tapestation (Agilent) and Qubit. Library preparation was performed using Ovation Solo RNA seq kit (Tecan, Switzerland) following manufacturer instructions. Library quality was assessed on tapestation (Agilent). Library pooling was done using MiSeq and pooled library were shipped to Novogene for sequencing on Illumina NovaSeq6000. Each sample contains at least >25 million total paired end reads of length 150 bp. Sequencing data was processed using *Partek™ Flow™* software, v11.0. Read alignment to the mouse genome build mm10 was performed with STAR 2.7.8a (Dobin et al. 2013) and Partek**^®^** E/M model was used to assemble gene level expression data from filtered alignments. Differential gene expression analysis was conducted using DESeq2 (Love, Huber, and Anders 2014).

Genes with cutoff (FDR < 0.05 and FC > |1.5|) were denoted as significant differentially expressed genes. Pathway analysis was performed using Enrichr (Chen et al. 2013). Enrichments of gene sets were performed using GSEA (Subramanian et al. 2005). Quantitative overlap analysis was performed using Eulerr (Larsson et al. 2024).

### Phagocytosis assay

Phagocytosis assay kit (IgG FITC) kit was purchased through Cayman Chemicals. For each condition (Scramble, sgBicra#2, sgBicra#3, inhibitor treatment) 20,000 cells were seeded overnight. The inhibitor treatment was done for two hours prior to addition of the FITC labeled beads as per the manufacturer protocol (1:200). Propidium Iodide (Thermo Fisher Scientific) was used as a counter stain as a marker for live/dead cells. Uptake of beads was quantified using flow cytometric analysis using Fortessa (BD Biosciences). Data were analyzed by FlowJo software (Tree Star, Ashland, OR).

### Hematocrit analysis

Peripheral blood was collected from 6-month-old C57BL/6J (CD45.2^+^) for comprehensive complete blood count analysis and processed for hematological analysis by IDEXX BioAnalytics, West Sacramento, CA.

### Statistical analyses and data availability

All statistical analyses and graphical visualizations were performed using Graphpad Prism9 software (GraphPad Software Inc). The data generated in this study are publicly available in Gene Expression Omnibus (GEO) upon publication. All other data generated in this study are available within the article and its supplementary data files.

## Acknowledgements

This work was supported by grants from the NIH (U01CA207532), the DoD (PCRP PC210004) and the Elsa U. Pardee Foundation to ECD. The authors gratefully acknowledge the support of the Purdue Computational Genomics Shared Resource and the Purdue Transgenic Mouse Facility and Flow Cytometry Facility from the Institute for Cancer Research, NIH grant P30 CA023168. The Flow Cytometry Facility also acknowledges Bindley Biosciences and NIH S10 ODO20029. We also acknowledge the support of the Collaborative Core for Cancer Bioinformatics from the Purdue University Institute for Cancer Research (PICR), NIH grants R01AI141439, P30CA023168 and the Walther Cancer Foundation. S.S. was supported by the Ross-Lynn Research Scholar Fund, Yeshwant A. Gore Graduate Grant from the Purdue University Institute for Cancer Research, and the Bilsland Dissertation Fellowship from PULSe. AA was supported by a Fullbright fellowship, the Lilly Graduate Fellowship from the Purdue College of Pharmacy, the SIRG grant administered through the Purdue Institute for Cancer Research, and the Bilsland Dissertation Fellowship from PULSe. GJ was supported by the SIRG grant administered through the Purdue Institute for Cancer Research. The authors would like to acknowledge Dr. Julie Lessard, Dr. Stephen Konieczny, Dr. Harm HogenEsch, Dr. Diana Hargreaves, and Dr. Helen McRae for their helpful advice and assistance and Dr. Brittany Allen-Petersen for microscopy help. Schematics were designed using Biorender.

---

**Supplementary Figure 1:**
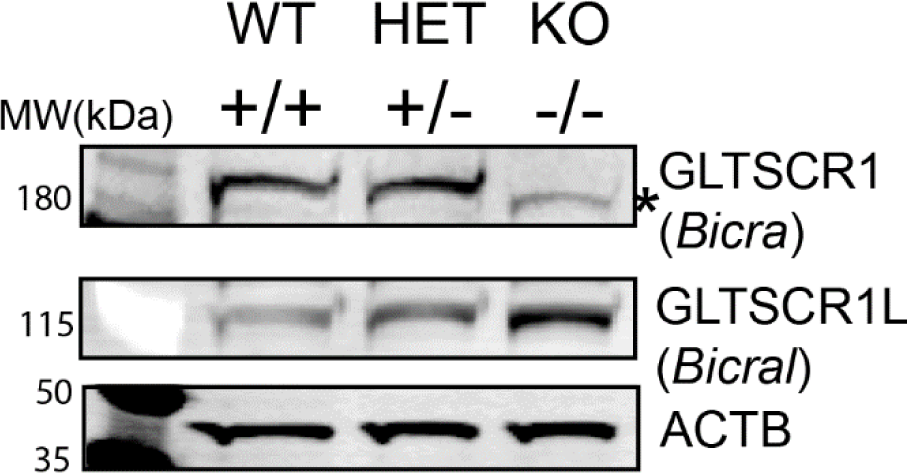
(A) Western blot performed from protein lysates isolated from mouse embryonic fibroblasts cultured *in vitro* from E12.5 embryos. *Shows a minor band that corresponds to GLTSCR1 isoform 2 with an alternate start site in the middle of exon 6 (Smith et al. 2000).

**Supplementary Figure 2:**
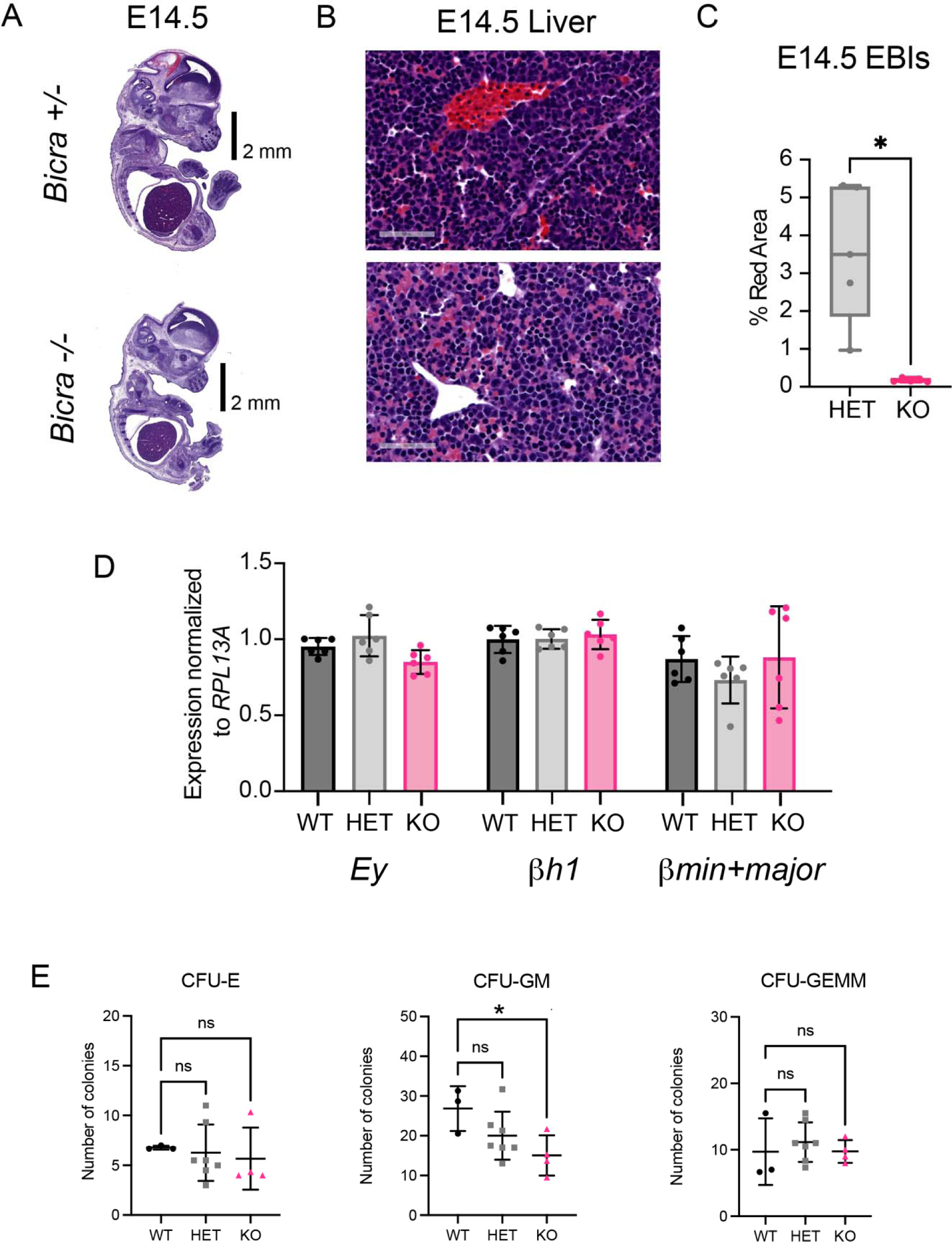
(A) Representative image of whole embryo H&E at E14.5 for *Bicra*^-/-^ pup and control littermates (B) Representative image for erythroblastic island (EBI) in fetal liver at E14.5 for *Bicra*^-/-^ knockout pup and control *Bicra*^-/+^ heterozygous littermates (C) Quantification of EBI in fetal liver at E14.5 birth for *Bicra*^-/-^ knockout pup and control *Bicra*^-/+^ heterozygous littermates using Image J software, n = 1 independent biological replicate experiment with 5 representative images/replicate taken for quantification, statistical test performed using Prism software Welch test (D) Quantification of colony forming units. Abbreviation: CFU-granulocyte, erythrocyte, monocyte/macrophage, megakaryocyte (GEMM); CFU-granulocyte-monocyte/macrophage (GM); E, CFU-erythroid, n = 3, n = 7 and n= 4 independent biological replicates for wildtype, heterozygous and homozygous genotype respectively. Mean summary statistics for each biological replicate are shown, Statistical testing is performed using Prism software with using Ordinary 1-way ANOVA (with multiple comparison). (E) Relative expression of ß globin genes by qRT-PCR, statistical analysis is performed using Prism software with 2way ANOVA (with multiple comparison). A designation of * = P ≤ 0.05.

**Supplementary Figure 3:**
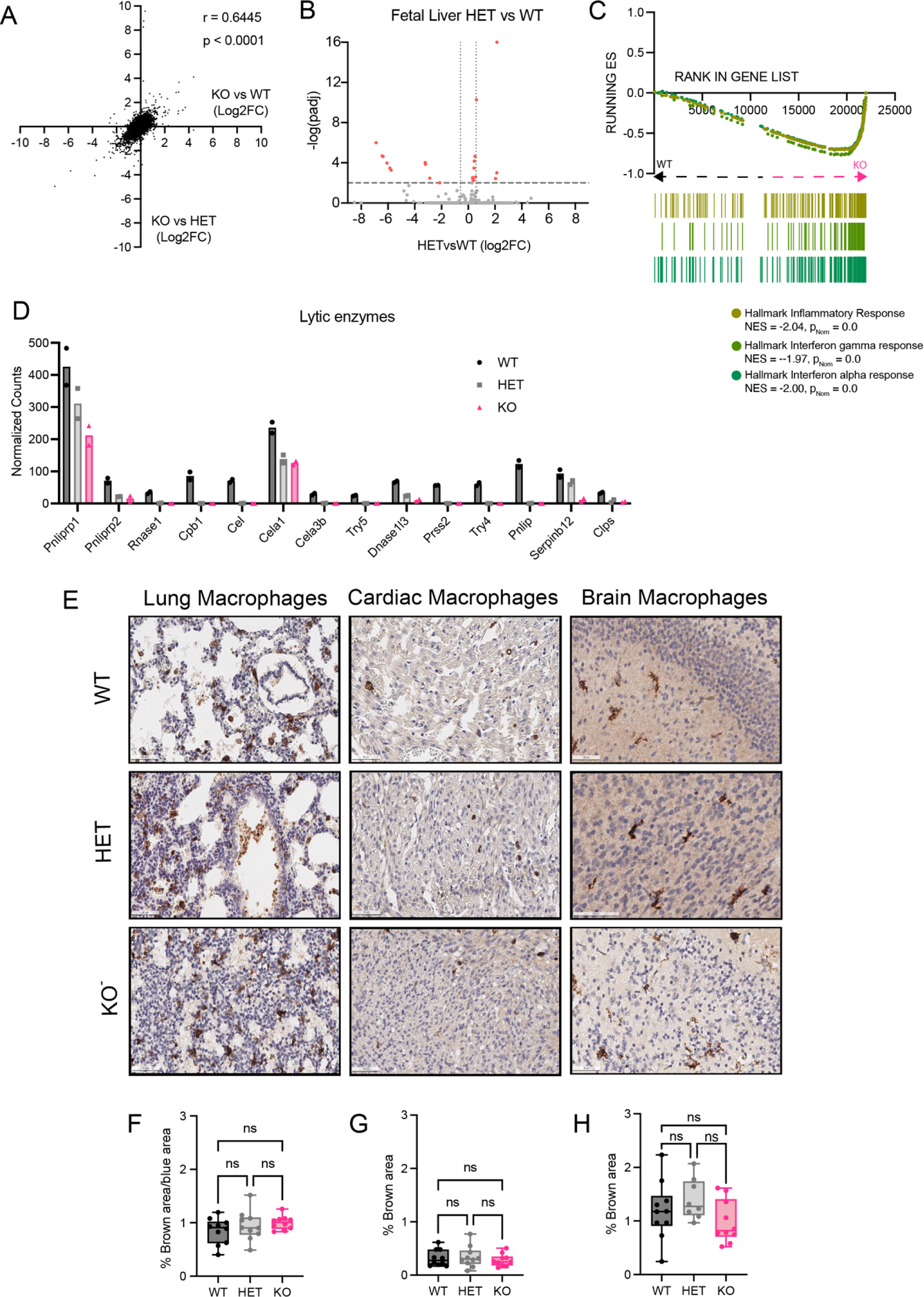
(A) Correlation of all gene expression changes in fetal liver bulk RNA-Seq for homozygous *Bicra*^-/-^ knockout mice compared to *Bicra*^+/+^ wildtype (*x*-axis) and *Bicra*^+/-^ heterozygous mice verses *Bicra*^+/+^ wildtype (*y*-axis). (B) Volcano plot of fetal liver bulk RNA-Seq gene expression changes in *Bicra*^+/-^ heterozygous compared to *Bicra*^+/+^ wildtype littermate controls at E14.5. The Log2FC indicates the mean expression level for each gene. Each dot represents one gene. The differentially expressed genes are shown in red with *p*_adj_ < 0.05 and FC > |1.5|. (C) GSEA analysis of RNA-Seq of bulk fetal livers from homozygous *Bicra*^-/-^ knockout mice compared to *Bicra*^+/+^ wildtype littermates. The most enriched Hallmark gene sets are depicted. (D) Lytic enzyme gene normalized counts between mice of different genotypes. (E) Representative image of F4/80 macrophage marker specific staining at E14.5 in lung, cardiac and brain tissue respectively for *Bicra*^-/-^ pups and control littermates. (F) Quantification of F4/80^+^ staining at E14.5 lung tissue using ImageJ software, n = 2 independent biological replicate experiments with 5 representative images/replicate taken for quantification, statistical test performed using Prism software using Ordinary 1-way ANOVA (with multiple comparisons). (G) Quantification of F4/80^+^ staining at E14.5 cardiac tissue section using ImageJ software, n = 2 independent biological replicate experiments with 5 representative images/replicate taken for quantification, statistical test performed using Prism software using Ordinary 1-way ANOVA (with multiple comparison). (H) Quantification of F4/80^+^ staining at E14.5 brain tissue section using ImageJ software, n = 2 independent biological replicate experiments with 5 representative images/replicate taken for quantification, statistical test performed using Prism software using Ordinary 1-way ANOVA (with multiple comparison). A designation of * = P ≤ 0.05; ** = P ≤ 0.01; *** = P ≤ 0.001; ns = non-significant (P > 0.05).

**Supplementary Figure 4:**
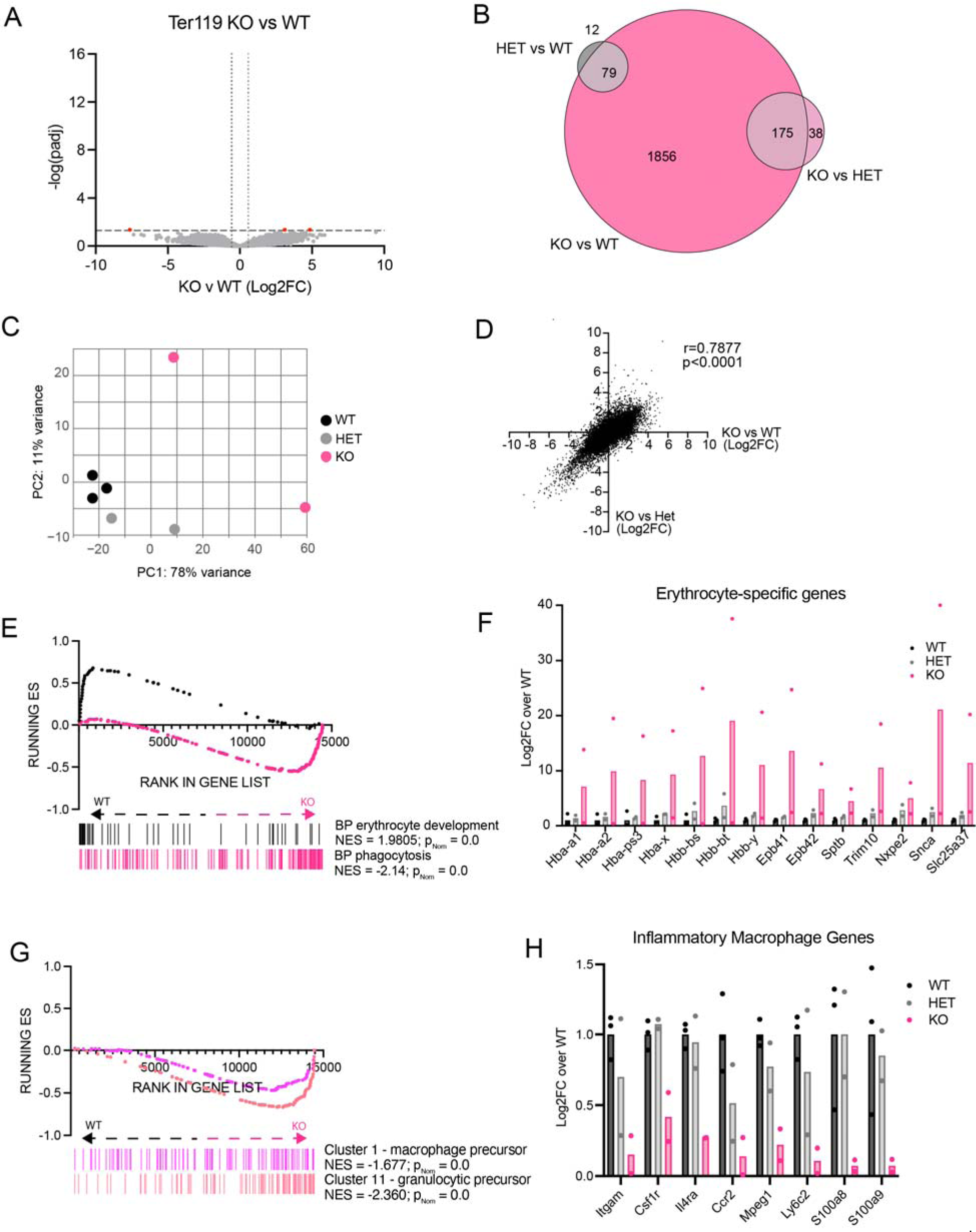
(A) Volcano plot of gene expression changes in Ter119^+^ sorted cells from *Bicra^-/-^* homozygous knockout compared to *Bicra^-+/+^*wildtype littermate controls at E14.5. The Log2 fold change indicates the mean expression level for each gene. Each dot represents one gene. The differentially expressed genes are shown in red with FDR < 0.05 and FC > |1.5|. (B) Overlap of differentially expressed genes (FDR < 0.05, FC > |1.5|) in F4/80^+^ sorted macrophages from *Bicra^-/-^*knockout mice compared to *Bicra^-+/+^* wildtype (bright pink), *Bicra^-/-^* knockout mice compared to *Bicra^-/+^*heterozygous mice (light pink) and *Bicra^-/+^* heterozygous mice compared to *Bicra^-+/+^* wildtype (grey). (C) Principal component analysis of F4/80^+^ sorted macrophages RNA-Seq samples isolated from fetal liver at E14.5. (D) Correlation of all gene expression changes in F4/80^+^ sorted macrophages RNA-Seq samples for homozygous knockout mice versus wildtype (*x*-axis) and heterozygous knockout mice verses wildtype (*y*-axis). (E) GSEA of RNA-Seq from fetal liver F4/80^+^ sorted macrophages from *Bicra^-+/+^* wildtype and *Bicra^-/-^* knockout mice against gene sets representing biological processes (BP). (F) Erythrocyte specific gene expression quantified using a comparison of normalized counts between F4/80^+^ sorted macrophages from mice of different genotypes. (G) GSEA of RNA-Seq from fetal liver F4/80^+^ sorted macrophages from *Bicra^-+/+^*wildtype and *Bicra^-/-^* knockout mice. Gene sets were created from the top 100 unique genes from each cluster defined using scRNA-Seq data of mouse fetal (E14.5) liver macrophages (Kayvanjoo et al. 2023). (H) Macrophage-specific gene expression depicted as the fold change (FC) of the average of the normalized counts for *Bicra^-+/+^* wildtype samples.

**Supplementary Figure 5:**
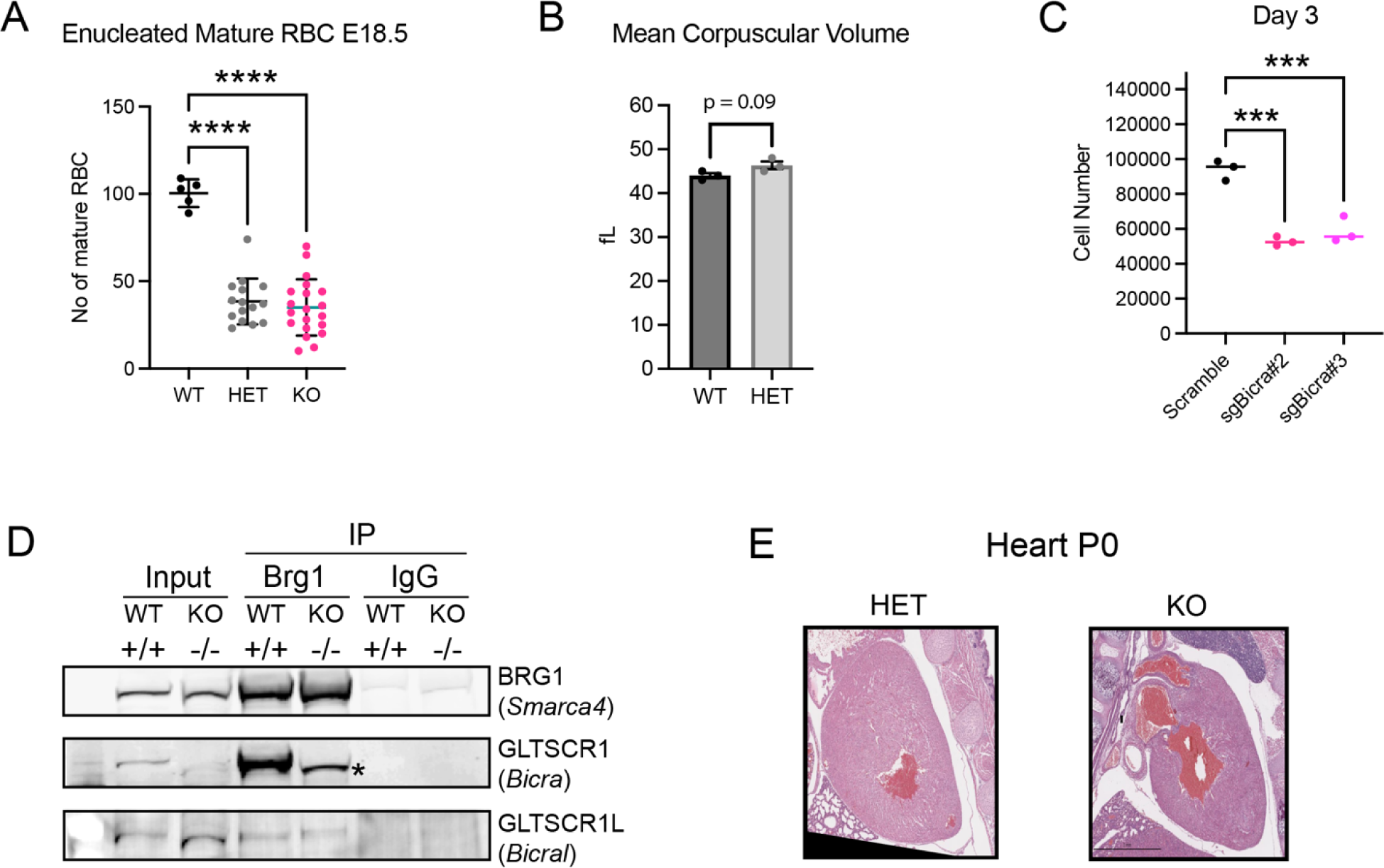
(A) Quantification of enucleated red blood cells in fetal liver H&E at E18.5 for *Bicra*^-/-^ knockout pup and control littermates, n = 1, n = 3, n = 4 independent biological replicates for wildtype, heterozygous and homozygous genotype, respectively; for each pup five fields of views are quantified, statistical test performed using Prism software using Ordinary 1-way ANOVA (with multiple comparison). (B) Quantification of mean corpuscular volume in peripheral blood of adult mice for *Bicra*^+/-^ heterozygous pup and wildtype control littermates using hematocrit analysis. (C) Quantification of cell number for the knockout lines compared to scramble upon 3 days of *Bicra* knockout in RAW 264.7 cell line; Statistical testing is performed using Prism software with Ordinary 1-way ANOVA (with multiple comparison). A designation of * = P ≤ 0.05; ** = P ≤ 0.01; *** = P ≤ 0.001; ns = non-significant (P > 0.05). (D) Immunoprecipitation with antibody against BRG1 (*Smarca4*) performed from protein lysates isolated from mouse embryonic fibroblasts cultured *in vitro* from E12.5 embryos. * Shows a minor band that corresponds to GLTSCR1 isoform 2 with an alternate start site in the middle of exon 6 (Smith et al. 2000). (E) Heart morphology at birth for *Bicra*^-/-^ knockout pup and control heterozygous littermates.

## Notes

### Competing Interest Statement

The authors have declared no competing interest.

## References

Ahmed, Nasiha S., Jovylyn Gatchalian, Josephine Ho, Mannix J. Burns, Nasun Hah, Zong Wei, Michael Downes, Ronald M. Evans, and Diana C. Hargreaves. 2022. “BRD9 Regulates Interferon-Stimulated Genes during Macrophage Activation via Cooperation with BET Protein BRD4.” Proceedings of the National Academy of Sciences 119 (1): e2110812119. 10.1073/pnas.2110812119.

Alpsoy, Aktan, and Emily C. Dykhuizen. 2018. “Glioma Tumor Suppressor Candidate Region Gene 1 (GLTSCR1) and Its Paralog GLTSCR1-like Form SWI/SNF Chromatin Remodeling Subcomplexes.” The Journal of Biological Chemistry 293 (11): 3892–3903. 10.1074/jbc.RA117.001065.

Alpsoy, Aktan, Sagar M. Utturkar, Benjamin C. Carter, Alisha Dhiman, Sandra E. Torregrosa-Allen, Melanie P. Currie, Bennett D. Elzey, and Emily C. Dykhuizen. 2021. “BRD9 Is a Critical Regulator of Androgen Receptor Signaling and Prostate Cancer Progression.” Cancer Research 81 (4): 820–33. 10.1158/0008-5472.CAN-20-1417.

Baggiolini, Marco, and Jörg Schnyder. 1982. “Synthesis and Release of Lytic Enzymes by Macrophages in Chronic Inflammation.” In Macrophages and Natural Killer Cells: Regulation and Function, edited by Sigurd J. Normann and Ernst Sorkin, 305–12. Boston, MA: Springer US. 10.1007/978-1-4684-4394-3_31.

Baron, Margaret H., Joan Isern, and Stuart T. Fraser. 2012. “The Embryonic Origins of Erythropoiesis in Mammals.” Blood 119 (21): 4828–37. 10.1182/blood-2012-01-153486.

Basu, Priyadarshi, Pamela E. Morris, Jack L. Haar, Maqsood A. Wani, Jerry B. Lingrel, Karin M. L. Gaensler, and Joyce A. Lloyd. 2005. “KLF2 Is Essential for Primitive Erythropoiesis and Regulates the Human and Murine Embryonic β-like Globin Genes in Vivo.” Blood 106 (7): 2566–71. 10.1182/blood-2005-02-0674.

Berger, Shelley L. 2007. “The Complex Language of Chromatin Regulation during Transcription.” Nature 447 (7143): 407–12. 10.1038/nature05915.

Bickmore, Wendy A., and Bas van Steensel. 2013. “Genome Architecture: Domain Organization of Interphase Chromosomes.” Cell 152 (6): 1270–84. 10.1016/j.cell.2013.02.001.

Bluemn, Theresa, Jesse Schmitz, Yuhong Chen, Yongwei Zheng, Yongguang Zhang, Shikan Zheng, Robert Burns, et al. 2021. “Arid2 Regulates Hematopoietic Stem Cell Differentiation in Normal Hematopoiesis.” Experimental Hematology 94 (February):37–46. 10.1016/j.exphem.2020.12.004.

Brien, Gerard L, David Remillard, Junwei Shi, Matthew L Hemming, Jonathon Chabon, Kieran Wynne, Eugène T Dillon, et al. 2018. “Targeted Degradation of BRD9 Reverses Oncogenic Gene Expression in Synovial Sarcoma.” Edited by Maarten van Lohuizen and Charles L Sawyers. eLife 7 (November):e41305. 10.7554/eLife.41305.

Bultman, Scott J., Thomas C. Gebuhr, and Terry Magnuson. 2005. “A Brg1 Mutation That Uncouples ATPase Activity from Chromatin Remodeling Reveals an Essential Role for SWI/SNF-Related Complexes in Beta-Globin Expression and Erythroid Development.” Genes & Development 19 (23): 2849–61. 10.1101/gad.1364105.

Cairns, Bradley R. 2007. “Chromatin Remodeling: Insights and Intrigue from Single-Molecule Studies.” Nature Structural & Molecular Biology 14 (11): 989–96. 10.1038/nsmb1333.

Chambers, Courtney, Katerina Cermakova, Yuen San Chan, Kristen Kurtz, Katharina Wohlan, Andrew Henry Lewis, Christiana Wang, et al. 2023. “SWI/SNF Blockade Disrupts PU.1-Directed Enhancer Programs in Normal Hematopoietic Cells and Acute Myeloid Leukemia.” Cancer Research 83 (7): 983–96. 10.1158/0008-5472.CAN-22-2129.

Chen, Edward Y., Christopher M. Tan, Yan Kou, Qiaonan Duan, Zichen Wang, Gabriela Vaz Meirelles, Neil R. Clark, and Avi Ma’ayan. 2013. “Enrichr: Interactive and Collaborative HTML5 Gene List Enrichment Analysis Tool.” BMC Bioinformatics 14 (April):128. 10.1186/1471-2105-14-128.

Chow, Andrew, Matthew Huggins, Jalal Ahmed, Daigo Hashimoto, Daniel Lucas, Yuya Kunisaki, Sandra Pinho, et al. 2013. “CD169+ Macrophages Provide a Niche Promoting Erythropoiesis under Homeostasis and Stress.” Nature Medicine 19 (4): 429–36. 10.1038/nm.3057.

Del Gaudio, Nunzio, Antonella Di Costanzo, Ning Qing Liu, Lidio Conte, Antimo Migliaccio, Michiel Vermeulen, Joost H. A. Martens, Hendrik G. Stunnenberg, Angela Nebbioso, and Lucia Altucci. 2019. “BRD9 Binds Cell Type-Specific Chromatin Regions Regulating Leukemic Cell Survival via STAT5 Inhibition.” Cell Death & Disease 10 (5): 1–14. 10.1038/s41419-019-1570-9.

Dickinson, Mary E., Ann M. Flenniken, Xiao Ji, Lydia Teboul, Michael D. Wong, Jacqueline K. White, Terrence F. Meehan, et al. 2016. “High-Throughput Discovery of Novel Developmental Phenotypes.” Nature 537 (7621): 508–14. 10.1038/nature19356.

Dobin, Alexander, Carrie A. Davis, Felix Schlesinger, Jorg Drenkow, Chris Zaleski, Sonali Jha, Philippe Batut, Mark Chaisson, and Thomas R. Gingeras. 2013. “STAR: Ultrafast Universal RNA-Seq Aligner.” Bioinformatics (Oxford, England) 29 (1): 15–21. 10.1093/bioinformatics/bts635.

Elrick, Hillary, Kevin A. Peterson, Joshua A. Wood, Denise G. Lanza, Elif F. Acar, Lydia Teboul, Edward J. Ryder, et al. 2021. “The Production of 4,182 Mouse Lines Identifies Experimental and Biological Variables Impacting Cas9-Mediated Mutant Mouse Line Production.” bioRxiv. 10.1101/2021.10.06.463037.

Frederick, Megan A., Kaylyn E. Williamson, Meilin Fernandez Garcia, Max B. Ferretti, Ryan L. McCarthy, Greg Donahue, Edgar Luzete Monteiro, et al. 2023. “A Pioneer Factor Locally Opens Compacted Chromatin to Enable Targeted ATP-Dependent Nucleosome Remodeling.” Nature Structural & Molecular Biology 30 (1): 31–37. 10.1038/s41594-022-00886-5.

Garg, Swati, Wei Ni, Basudev Chowdhury, Ellen L. Weisberg, Martin Sattler, and James D. Griffin. 2024. “BRD9 Regulates Normal Human Hematopoietic Stem Cell Function and Lineage Differentiation.” Cell Death & Differentiation, May, 1–13. 10.1038/s41418-024-01306-5.

Gatchalian, Jovylyn, Shivani Malik, Josephine Ho, Dong-Sung Lee, Timothy W. R. Kelso, Maxim N. Shokhirev, Jesse R. Dixon, and Diana C. Hargreaves. 2018. “A Non-Canonical BRD9-Containing BAF Chromatin Remodeling Complex Regulates Naive Pluripotency in Mouse Embryonic Stem Cells.” Nature Communications 9 (1): 5139. 10.1038/s41467-018-07528-9.

Goonesekere, Nalin C. W., Wyatt Andersen, Alex Smith, and Xiaosheng Wang. 2018. “Identification of Genes Highly Downregulated in Pancreatic Cancer through a Meta-Analysis of Microarray Datasets: Implications for Discovery of Novel Tumor-Suppressor Genes and Therapeutic Targets.” Journal of Cancer Research and Clinical Oncology 144 (2): 309–20. 10.1007/s00432-017-2558-4.

Graf, Thomas, and Tariq Enver. 2009. “Forcing Cells to Change Lineages.” Nature 462 (7273): 587–94. 10.1038/nature08533.

Groza, Tudor, Federico Lopez Gomez, Hamed Haseli Mashhadi, Violeta Muñoz-Fuentes, Osman Gunes, Robert Wilson, Pilar Cacheiro, et al. 2023. “The International Mouse Phenotyping Consortium: Comprehensive Knockout Phenotyping Underpinning the Study of Human Disease.” Nucleic Acids Research 51 (D1): D1038–45. 10.1093/nar/gkac972.

Gutiérrez, José, Roberto Paredes, Fernando Cruzat, David A. Hill, Andre J. van Wijnen, Jane B. Lian, Gary S. Stein, Janet L. Stein, Anthony N. Imbalzano, and Martin Montecino. 2007. “Chromatin Remodeling by SWI/SNF Results in Nucleosome Mobilization to Preferential Positions in the Rat Osteocalcin Gene Promoter*.” Journal of Biological Chemistry 282 (13): 9445–57. 10.1074/jbc.M609847200.

Han, Fengyan, Beibei Yang, Yan Chen, Lu Liu, Xiaoqing Cheng, Jiaqi Huang, Ke Zhou, et al. 2023. “Loss of GLTSCR1 Causes Congenital Heart Defects by Regulating NPPA Transcription.” Angiogenesis 26 (2): 217–32. 10.1007/s10456-023-09869-6.

Han, Lin, Vikas Madan, Anand Mayakonda, Pushkar Dakle, Teoh Weoi Woon, Pavithra Shyamsunder, Hazimah Binte Mohd Nordin, et al. 2019. “Chromatin Remodeling Mediated by ARID1A Is Indispensable for Normal Hematopoiesis in Mice.” Leukemia 33 (9): 2291–2305. 10.1038/s41375-019-0438-4.

Hanspal, Manjit, and Jatinder S. Hanspal. 1994. “The Association of Erythroblasts With Macrophages Promotes Erythroid Proliferation and Maturation: A 30-kD Heparin-Binding Protein Is Involved in This Contact.” Blood 84 (10): 3494–3504. 10.1182/blood.V84.10.3494.3494.

Hanspal, Manjit, Yva Smockova, and Quang Uong. 1998. “Molecular Identification and Functional Characterization of a Novel Protein That Mediates the Attachment of Erythroblasts to Macrophages.” Blood 92 (8): 2940–50. 10.1182/blood.V92.8.2940.

Ho, Lena, and Gerald R. Crabtree. 2010. “Chromatin Remodelling during Development.” Nature 463 (7280): 474–84. 10.1038/nature08911.

Hoeffel, Guillaume, Jinmiao Chen, Yonit Lavin, Donovan Low, Francisca F. Almeida, Peter See, Anna E. Beaudin, et al. 2015. “C-Myb+ Erythro-Myeloid Progenitor-Derived Fetal Monocytes Give Rise to Adult Tissue-Resident Macrophages.” Immunity 42 (4): 665–78. 10.1016/j.immuni.2015.03.011.

Imbalzano, Anthony N., Hyockman Kwon, Michael R. Green, and Robert E. Kingston. 1994. “Facilitated Binding of TATA-Binding Protein to Nucleosomal DNA.” Nature 370 (6489): 481–85. 10.1038/370481a0.

Inoue, Daichi, Guo-Liang Chew, Bo Liu, Brittany C. Michel, Joseph Pangallo, Andrew R. D’Avino, Tyler Hitchman, et al. 2019. “Spliceosomal Disruption of the Non-Canonical BAF Complex in Cancer.” Nature 574 (7778): 432–36. 10.1038/s41586-019-1646-9.

Iwasaki, Hiromi, Chamorro Somoza, Hirokazu Shigematsu, Estelle A. Duprez, Junko Iwasaki-Arai, Shin-Ichi Mizuno, Yojiro Arinobu, et al. 2005. “Distinctive and Indispensable Roles of PU.1 in Maintenance of Hematopoietic Stem Cells and Their Differentiation.” Blood 106 (5): 1590–1600. 10.1182/blood-2005-03-0860.

Kawane, K., H. Fukuyama, G. Kondoh, J. Takeda, Y. Ohsawa, Y. Uchiyama, and S. Nagata. 2001. “Requirement of DNase II for Definitive Erythropoiesis in the Mouse Fetal Liver.” Science (New York, N.Y.) 292 (5521): 1546–49. 10.1126/science.292.5521.1546.

Kayvanjoo, Amir Hossein, Iva Splichalova, David Alejandro Bejarano, Hao Huang, Katharina Mauel, Nikola Makdissi, David Heider, et al. 2024. “Fetal Liver Macrophages Contribute to the Hematopoietic Stem Cell Niche by Controlling Granulopoiesis.” Edited by Florent Ginhoux and Pramod K Mistry. eLife 13 (March):e86493. 10.7554/eLife.86493.

Kornberg, R. D. 1974. “Chromatin Structure: A Repeating Unit of Histones and DNA.” Science (New York, N.Y.) 184 (4139): 868–71. 10.1126/science.184.4139.868.

Koulnis, Miroslav, Ramona Pop, Ermelinda Porpiglia, Jeffrey R. Shearstone, Daniel Hidalgo, and Merav Socolovsky. 2011. “Identification and Analysis of Mouse Erythroid Progenitors Using the CD71/TER119 Flow-Cytometric Assay.” Journal of Visualized Experiments: JoVE, no. 54 (August), 2809. 10.3791/2809.

Krasteva, Veneta, Manuel Buscarlet, Abigail Diaz-Tellez, Marie-Anne Bernard, Gerald R. Crabtree, and Julie A. Lessard. 2012. “The BAF53a Subunit of SWI/SNF-like BAF Complexes Is Essential for Hemopoietic Stem Cell Function.” Blood 120 (24): 4720–32. 10.1182/blood-2012-04-427047.

Krasteva, Veneta, Gerald R. Crabtree, and Julie A. Lessard. 2017. “The BAF45a/PHF10 Subunit of SWI/SNF-like Chromatin Remodeling Complexes Is Essential for Hematopoietic Stem Cell Maintenance.” Experimental Hematology 48 (April):58–71.e15. 10.1016/j.exphem.2016.11.008.

Krosl, Jana, Aline Mamo, Jalila Chagraoui, Brian T. Wilhelm, Simon Girard, Isabelle Louis, Julie Lessard, Claude Perreault, and Guy Sauvageau. 2010. “A Mutant Allele of the Swi/Snf Member BAF250a Determines the Pool Size of Fetal Liver Hemopoietic Stem Cell Populations.” Blood 116 (10): 1678–84. 10.1182/blood-2010-03-273862.

Kurata, Keiji, Mehmet K. Samur, Priscilla Liow, Kenneth Wen, Leona Yamamoto, Jiye Liu, Eugenio Morelli, et al. 2023. “BRD9 Degradation Disrupts Ribosome Biogenesis in Multiple Myeloma.” Clinical Cancer Research 29 (9): 1807–21. 10.1158/1078-0432.CCR-22-3668.

Lara-Astiaso, David, Ainhoa Goñi-Salaverri, Julen Mendieta-Esteban, Nisha Narayan, Cynthia Del Valle, Torsten Gross, George Giotopoulos, et al. 2023a. “In Vivo Screening Characterizes Chromatin Factor Functions during Normal and Malignant Hematopoiesis.” Nature Genetics 55 (9): 1542–54. 10.1038/s41588-023-01471-2.

Lara-Astiaso, David, Ainhoa Goñi-Salaverri, Julen Mendieta-Esteban, Nisha Narayan, Cynthia Del Valle, Torsten Gross, George Giotopoulos, et al. 2023b. “In Vivo Screening Characterizes Chromatin Factor Functions during Normal and Malignant Hematopoiesis.” Nature Genetics, August, 1–13. 10.1038/s41588-023-01471-2.

Larsson, Johan A. Jonathan R. Godfrey, Peter Gustafsson, David H. Eberly (geometric algorithms), Emanuel Huber (root solver code), and Florian Privé. 2024. “Eulerr: Area-Proportional Euler and Venn Diagrams with Ellipses.” https://cran.r-project.org/web/packages/eulerr/index.html.

Lee, Gloria, Annie Lo, Sarah A. Short, Tosti J. Mankelow, Frances Spring, Stephen F. Parsons, Karina Yazdanbakhsh, Narla Mohandas, David J. Anstee, and Joel Anne Chasis. 2006. “Targeted Gene Deletion Demonstrates That the Cell Adhesion Molecule ICAM-4 Is Critical for Erythroblastic Island Formation.” Blood 108 (6): 2064–71. 10.1182/blood-2006-03-006759.

Lewis, Kyle, Momoko Yoshimoto, and Takanori Takebe. 2021. “Fetal Liver Hematopoiesis: From Development to Delivery.” Stem Cell Research & Therapy 12 (1): 139. 10.1186/s13287-021-02189-w.

Li, Wei, Rongqun Guo, Yongping Song, and Zhongxing Jiang. 2021. “Erythroblastic Island Macrophages Shape Normal Erythropoiesis and Drive Associated Disorders in Erythroid Hematopoietic Diseases.” Frontiers in Cell and Developmental Biology 8 (February):613885. 10.3389/fcell.2020.613885.

Li, Wei, Yaomei Wang, Huizhi Zhao, Huan Zhang, Yuanlin Xu, Shihui Wang, Xinhua Guo, et al. 2019. “Identification and Transcriptome Analysis of Erythroblastic Island Macrophages.” Blood 134 (5): 480–91. 10.1182/blood.2019000430.

Liu, Jing, Xinhua Guo, Narla Mohandas, Joel A. Chasis, and Xiuli An. 2010. “Membrane Remodeling during Reticulocyte Maturation.” Blood 115 (10): 2021–27. 10.1182/blood-2009-08-241182.

Liu, Lulu, Xiaoling Wan, Peipei Zhou, Xiaoyuan Zhou, Wei Zhang, Xinhui Hui, Xiujie Yuan, et al. 2018. “The Chromatin Remodeling Subunit Baf200 Promotes Normal Hematopoiesis and Inhibits Leukemogenesis.” Journal of Hematology & Oncology 11 (1): 27. 10.1186/s13045-018-0567-7.

Livak, K. J., and T. D. Schmittgen. 2001. “Analysis of Relative Gene Expression Data Using Real-Time Quantitative PCR and the 2(-Delta Delta C(T)) Method.” Methods (San Diego, Calif.) 25 (4): 402–8. 10.1006/meth.2001.1262.

Love, Michael I., Wolfgang Huber, and Simon Anders. 2014. “Moderated Estimation of Fold Change and Dispersion for RNA-Seq Data with DESeq2.” Genome Biology 15 (12): 550. 10.1186/s13059-014-0550-8.

Luger, K., A. W. Mäder, R. K. Richmond, D. F. Sargent, and T. J. Richmond. 1997. “Crystal Structure of the Nucleosome Core Particle at 2.8 A Resolution.” Nature 389 (6648): 251–60. 10.1038/38444.

Madan, Vikas, Pavithra Shyamsunder, Pushkar Dakle, Weoi Woon Teoh, Lin Han, Zeya Cao, Hazimah Mohd Nordin, et al. 2023. “Dissecting the Role of SWI/SNF Component ARID1B in Steady State Hematopoiesis.” Blood Advances, August, bloodadvances.2023009946. 10.1182/bloodadvances.2023009946.

Magae, Junji, Toshimi Nagi, Kazuaki Takaku, Takao Kataoka, Hiroyuki Koshino, Masakazu Uramoto, and Kazuo Nagai. 1994. “Screening for Specific Inhibitors of Phagocytosis of Thioglycollate-Elicited Macrophages.” Bioscience, Biotechnology, and Biochemistry 58 (1): 104–7. 10.1271/bbb.58.104.

Manwani, Deepa, and James J. Bieker. 2008. “Chapter 2 The Erythroblastic Island.” In Current Topics in Developmental Biology, 82:23–53. Red Cell Development. Academic Press. 10.1016/S0070-2153(07)00002-6.

Mas, Gloria, Na Man, Yuichiro Nakata, Concepcion Martinez-Caja, Daniel L. Karl, Felipe Beckedorff, Francesco Tamiro, et al. 2023. “The SWI/SNF Chromatin Remodeling Subunit DPF2 Facilitates NRF2-Dependent Anti-Inflammatory and Anti-Oxidant Gene Expression.” The Journal of Clinical Investigation, May, e158419. 10.1172/JCI158419.

Mashtalir, Nazar, Andrew R. D’Avino, Brittany C. Michel, Jie Luo, Joshua Pan, Jordan E. Otto, Hayley J. Zullow, et al. 2018. “Modular Organization and Assembly of SWI/SNF Family Chromatin Remodeling Complexes.” Cell 175 (5): 1272–1288.e20. 10.1016/j.cell.2018.09.032.

Michel, Brittany C., Andrew R. D’Avino, Seth H. Cassel, Nazar Mashtalir, Zachary M. McKenzie, Matthew J. McBride, Alfredo M. Valencia, et al. 2018. “A Non-Canonical SWI/SNF Complex Is a Synthetic Lethal Target in Cancers Driven by BAF Complex Perturbation.” Nature Cell Biology 20 (12): 1410–20. 10.1038/s41556-018-0221-1.

Mukherjee, Kaustav, and James J. Bieker. 2021. “Transcriptional Control of Gene Expression and the Heterogeneous Cellular Identity of Erythroblastic Island Macrophages.” Frontiers in Genetics 12 (November). 10.3389/fgene.2021.756028.

Mukherjee, Kaustav, Li Xue, Antanas Planutis, Merlin Nithya Gnanapragasam, Andrew Chess, and James J Bieker. 2021. “EKLF/KLF1 Expression Defines a Unique Macrophage Subset during Mouse Erythropoiesis.” Edited by Florent Ginhoux, Carla V Rothlin, and Christian Schulz. eLife 10 (February):e61070. 10.7554/eLife.61070.

Naidu, Samisubbu R., Maegan Capitano, James Ropa, Scott Cooper, Xinxin Huang, and Hal E. Broxmeyer. 2022. “Chromatin Remodeling Subunit BRM and Valine Regulate Hematopoietic Stem/Progenitor Cell Function and Self-Renewal via Intrinsic and Extrinsic Effects.” Leukemia 36 (3): 821–33. 10.1038/s41375-021-01426-8.

Orkin, Stuart H., and Leonard I. Zon. 2008. “Hematopoiesis: An Evolving Paradigm for Stem Cell Biology.” Cell 132 (4): 631–44. 10.1016/j.cell.2008.01.025.

Palis, James. 2017. “Interaction of the Macrophage and Primitive Erythroid Lineages in the Mammalian Embryo.” Frontiers in Immunology 7. https://www.frontiersin.org/articles/10.3389/fimmu.2016.00669.

Palis, James, Scott Robertson, Marion Kennedy, Charles Wall, and Gordon Keller. 1999. “Development of Erythroid and Myeloid Progenitors in the Yolk Sac and Embryo Proper of the Mouse.” Development 126 (22): 5073–84. 10.1242/dev.126.22.5073.

Popescu, Dorin-Mirel, Rachel A. Botting, Emily Stephenson, Kile Green, Simone Webb, Laura Jardine, Emily F. Calderbank, et al. 2019. “Decoding Human Fetal Liver Haematopoiesis.” Nature 574 (7778): 365–71. 10.1038/s41586-019-1652-y.

Porcu, Susanna, Maria F. Manchinu, Maria F. Marongiu, Valeria Sogos, Daniela Poddie, Isadora Asunis, Loredana Porcu, et al. 2011. “Klf1 Affects DNase II-Alpha Expression in the Central Macrophage of a Fetal Liver Erythroblastic Island: A Non-Cell-Autonomous Role in Definitive Erythropoiesis.” Molecular and Cellular Biology 31 (19): 4144–54. 10.1128/MCB.05532-11.

Raschke, W. C., S. Baird, P. Ralph, and I. Nakoinz. 1978. “Functional Macrophage Cell Lines Transformed by Abelson Leukemia Virus.” Cell 15 (1): 261–67. 10.1016/0092-8674(78)90101-0.

Romano, Laurel, Katie G. Seu, Julien Papoin, David E. Muench, Diamantis Konstantinidis, André Olsson, Katrina Schlum, et al. 2022. “Erythroblastic Islands Foster Granulopoiesis in Parallel to Terminal Erythropoiesis.” Blood 140 (14): 1621–34. 10.1182/blood.2022015724.

Sadahira, Y., T. Yoshino, and Y. Monobe. 1995. “Very Late Activation Antigen 4-Vascular Cell Adhesion Molecule 1 Interaction Is Involved in the Formation of Erythroblastic Islands.” Journal of Experimental Medicine 181 (1): 411–15. 10.1084/jem.181.1.411.

Seu, Katie Giger, Julien Papoin, Rose Fessler, Jimmy Hom, Gang Huang, Narla Mohandas, Lionel Blanc, and Theodosia A. Kalfa. 2017. “Unraveling Macrophage Heterogeneity in Erythroblastic Islands.” Frontiers in Immunology 8 (September). 10.3389/fimmu.2017.01140.

Skutelsky, Ehud, and David Danon. 1972. “On the Expulsion of the Erythroid Nucleus and Its Phagocytosis.” The Anatomical Record 173 (1): 123–26. 10.1002/ar.1091730111.

Smith, Justin S., Issei Tachibana, Ute Pohl, Hyun K. Lee, Uma Thanarajasingam, Bryce P. Portier, Keisuke Ueki, et al. 2000. “A Transcript Map of the Chromosome 19q-Arm Glioma Tumor Suppressor Region.” Genomics 64 (1): 44–50. 10.1006/geno.1999.6101.

Soni, Shivani, Shashi Bala, Babette Gwynn, Kenneth E. Sahr, Luanne L. Peters, and Manjit Hanspal. 2006. “Absence of Erythroblast Macrophage Protein (Emp) Leads to Failure of Erythroblast Nuclear Extrusion *.” Journal of Biological Chemistry 281 (29): 20181–89. 10.1074/jbc.M603226200.

Subramanian, Aravind, Pablo Tamayo, Vamsi K. Mootha, Sayan Mukherjee, Benjamin L. Ebert, Michael A. Gillette, Amanda Paulovich, et al. 2005. “Gene Set Enrichment Analysis: A Knowledge-Based Approach for Interpreting Genome-Wide Expression Profiles.” Proceedings of the National Academy of Sciences of the United States of America 102 (43): 15545–50. 10.1073/pnas.0506580102.

Tao, Hai-Ping, Teng-Fei Lu, Shuang Li, Gong-Xue Jia, Xiao-Na Zhang, Qi-En Yang, and Yun-Peng Hou. 2023. “Pancreatic Lipase-Related Protein 2 Is Selectively Expressed by Peritubular Myoid Cells in the Murine Testis and Sustains Long-Term Spermatogenesis.” Cellular and Molecular Life Sciences 80 (8): 217. 10.1007/s00018-023-04872-y.

Tu, Jiayi, Xiliang Liu, Haibo Jia, James Reilly, Shanshan Yu, Chen Cai, Fei Liu, et al. 2020. “The Chromatin Remodeler Brg1 Is Required for Formation and Maintenance of Hematopoietic Stem Cells.” FASEB Journal: Official Publication of the Federation of American Societies for Experimental Biology 34 (9): 11997–8. 10.1096/fj.201903168RR.

Tu, Zhaowei, and Yi Zheng. 2022. “Role of ATP-Dependent Chromatin Remodelers in Hematopoietic Stem and Progenitor Cell Maintenance.” Current Opinion in Hematology 29 (4): 174–80. 10.1097/MOH.0000000000000710.

Wang, Liu, Tae Gyu Oh, Jason Magida, Gabriela Estepa, S. M. Bukola Obayomi, Ling-Wa Chong, Jovylyn Gatchalian, et al. 2021. “Bromodomain Containing 9 (BRD9) Regulates Macrophage Inflammatory Responses by Potentiating Glucocorticoid Receptor Activity.” Proceedings of the National Academy of Sciences 118 (35): e2109517118. 10.1073/pnas.2109517118.

Wang, W., J. Côté, Y. Xue, S. Zhou, P. A. Khavari, S. R. Biggar, C. Muchardt, et al. 1996. “Purification and Biochemical Heterogeneity of the Mammalian SWI-SNF Complex.” The EMBO Journal 15 (19): 5370–82.

Wang, Xiaofeng, Su Wang, Emma C. Troisi, Thomas P. Howard, Jeffrey R. Haswell, Bennett K. Wolf, William H. Hawk, et al. 2019. “BRD9 Defines a SWI/SNF Sub-Complex and Constitutes a Specific Vulnerability in Malignant Rhabdoid Tumors.” Nature Communications 10 (1): 1881. 10.1038/s41467-019-09891-7.

Weisberg, Ellen, Basudev Chowdhury, Chengcheng Meng, Abigail E. Case, Wei Ni, Swati Garg, Martin Sattler, et al. 2022. “BRD9 Degraders as Chemosensitizers in Acute Leukemia and Multiple Myeloma.” Blood Cancer Journal 12 (7): 1–10. 10.1038/s41408-022-00704-7.

Weissman, I. L. 2000. “Stem Cells: Units of Development, Units of Regeneration, and Units in Evolution.” Cell 100 (1): 157–68. 10.1016/s0092-8674(00)81692-x.

Woodcock, Christopher L., and Rajarshi P. Ghosh. 2010. “Chromatin Higher-Order Structure and Dynamics.” Cold Spring Harbor Perspectives in Biology 2 (5): a000596. 10.1101/cshperspect.a000596.

Wu, Jun, Karen Krchma, Hyung Joo Lee, Sairam Prabhakar, Xiaoli Wang, Haiyong Zhao, Xiaoyun Xing, et al. 2020. “Requisite Chromatin Remodeling for Myeloid and Erythroid Lineage Differentiation from Erythromyeloid Progenitors.” Cell Reports 33 (7): 108395. 10.1016/j.celrep.2020.108395.

Xiao, Muran, Shinji Kondo, Masaki Nomura, Shinichiro Kato, Koutarou Nishimura, Weijia Zang, Yifan Zhang, et al. 2023. “BRD9 Determines the Cell Fate of Hematopoietic Stem Cells by Regulating Chromatin State.” Nature Communications 14 (December):8372. 10.1038/s41467-023-44081-6.

Zhao, Xinhui, Boris Bartholdy, Yukiya Yamamoto, Erica K. Evans, Meritxell Alberich-Jordà, Philipp B. Staber, Touati Benoukraf, et al. 2022. “PU.1-c-Jun Interaction Is Crucial for PU.1 Function in Myeloid Development.” Communications Biology 5 (1): 1–15. 10.1038/s42003-022-03888-7.

